# An EMT-primary cilium-GLIS2 signaling axis regulates mammogenesis and claudin-low breast tumorigenesis

**DOI:** 10.1101/2020.12.29.424695

**Authors:** Molly M. Wilson, Céline Callens, Matthieu Le Gallo, Svetlana Mironov, Qiong Ding, Amandine Salamagnon, Tony E. Chavarria, Abena D. Peasah, Arjun Bhutkar, Sophie Martin, Florence Godey, Patrick Tas, Anton M. Jetten, Jane E. Visvader, Robert A. Weinberg, Massimo Attanasio, Claude Prigent, Jacqueline A. Lees, Vincent J Guen

## Abstract

The Epithelial–Mesenchymal Transition (EMT) and primary ciliogenesis induce stem cell properties in basal Mammary Stem Cells (MaSCs) to promote mammogenesis, but the underlying mechanisms remain incompletely understood. Here, we show that EMT transcription factors promote ciliogenesis at intermediate EMT transition states by activating ciliogenesis inducers, including FGFR1. The resulting primary cilia promote BBS11-dependent ubiquitination and inactivation of a central signaling node, GLIS2. We show that GLIS2 inactivation promotes MaSC stemness, and GLIS2 is required for normal mammary gland development. Moreover, GLIS2 inactivation is required to induce the proliferative and tumorigenic capacities of the Mammary-Tumor-initiating cells (MaTICs) of claudin-low breast cancers. Claudin-low breast tumors can be segregated from other breast tumor subtypes based on a GLIS2-dependent gene expression signature. Collectively, our findings establish molecular mechanisms by which EMT programs induce ciliogenesis to control MaSC and MaTIC biology, mammary gland development, and claudin-low breast cancer formation.

## Introduction

The mammary gland is a branched ductal tissue comprised of a stratified epithelium containing luminal and basal cells surrounded by a basement membrane embedded in stroma(*1*). Mammary gland development is fueled by basal multipotent and lineage-restricted unipotent mammary stem cells (MaSCs) (*1*). The stem cell properties of basal MaSCs are induced by Epithelial-Mesenchymal transitions (EMT), a cell-biological program that enables epithelial cells to acquire an array of mesenchymal phenotypes (*2-5*). The EMT transcription factor (EMT-TF) Slug is expressed in MaSC-enriched basal cells (*2-4, 6, 7*). The ability of these stem cells to form organoids and reconstitute the mammary epithelium in transplantation experiments is enhanced or impaired by Slug overexpression or loss, respectively (*3, 4, 7*). Consistent with these findings, Slug-knockout (KO) mice demonstrate delayed mammary gland development (*6*).

Breast cancer comprises a variety of tumor types that are classified into therapeutic and molecular subtypes (*8*). The Triple-Negative Breast Cancer (TNBC) therapeutic subtype, in particular, is composed of breast tumors that do not express nuclear hormone receptors and HER2 and includes the less differentiated claudin-low molecular subtype (*8*). Claudin-low breast tumors are breast cancers characterized by low expression of epithelial cell-cell adhesion genes and high expression of EMT and basal MaSC genes (*8*). In murine carcinoma models of claudin-low breast cancer, the tumorigenic activity of Mammary Tumor-Initiating Cells (MaTICs) relies on EMT-TFs (*2, 3, 9-11*).

Collectively, these studies revealed that EMT programs tightly regulate mammary gland development and tumorigenesis by controlling both MaSCs and MaTICs. Recent work revealed that EMT programs promote MaSC and MaTIC stemness through induction of primary ciliogenesis (*11*). Primary ciliogenesis is a dynamic process in which a single microtubule-based structure, the primary cilium, is assembled by a modified version of the mother centriole, called the basal body, at the plasma membrane and protrudes from the surface, where it acts as a cell signaling center (*12*). Primary ciliogenesis requires the coordination of many cellular processes including a motor-driven process that involves bidirectional movement of multiprotein complexes along the cilium, called intraflagellar transport (IFT) (*12*). In the mammary gland, the molecular mechanisms linking EMT, primary cilia, and stemness are largely unknown.

Here, we show that EMT programs induce primary ciliogenesis at intermediate epithelial-mesenchymal transition states in both human and mouse mammary glands. Additionally, we establish that EMT-TFs induce the expression of direct ciliogenesis regulators and IFT inducers, including FGFR1, to promote primary ciliogenesis. We show that primary cilia control BBS11-dependent ubiquitination, and inactivation of a central signaling node, GLIS2. Furthermore, we demonstrate that GLIS2 inactivation promotes the stemness of MaSCs and MaTICs, mammogenesis and claudin-low breast tumorigenesis.

## Results

### EMT programs induce primary ciliogenesis at intermediate transition states in the mammary gland

We previously established that EMT programs are associated with primary ciliogenesis in the mouse mammary gland (*11*), spurred by our discovery that primary cilia were present in a high fraction of basal Slug-expressing cells present in ducts and terminal end buds in the murine mammary epithelium (Fig. S1A) (*11*). To determine whether this was also true in the human gland, we analyzed sections of human reduction mammoplasty tissue. Cells expressing the EMT-TF Slug were detected in the basal layer of the human mammary epithelium and expressed significantly lower levels of the epithelial marker E-cadherin than did the Slug-negative luminal cells (Fig. 1A). Importantly, primary cilia (detected by the cilium marker Arl13b and the centrosome marker γTubulin) marked the vast majority of Slug-expressing cells, in contrast to the Slug-negative cells (Fig. 1A). Hence, basal mammary epithelial cells, which exist in an intermediate EMT transition state, assemble primary cilia in both human and mouse mammary glands.

**Fig. 1:**
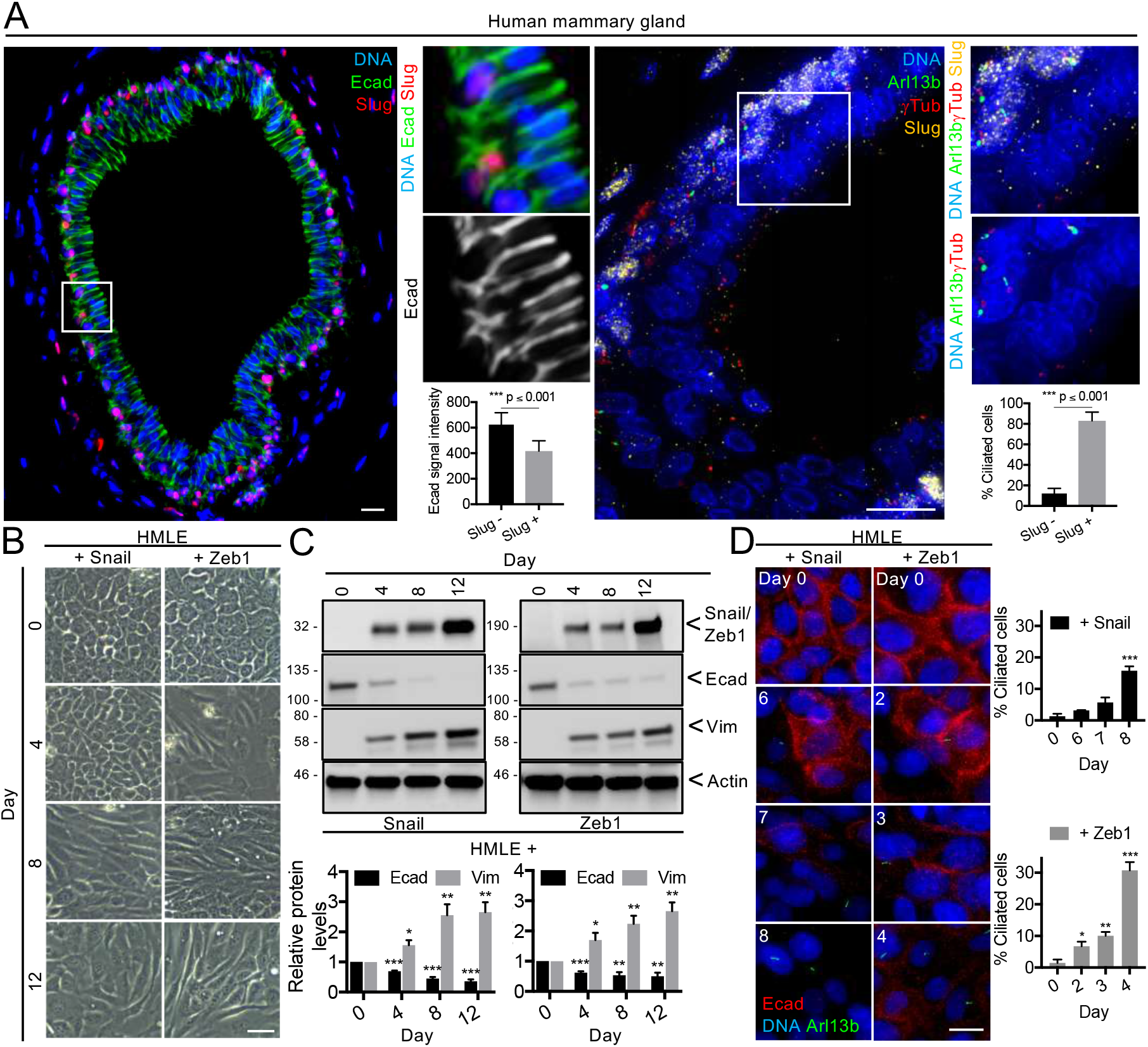
EMT programs induce primary ciliogenesis at intermediate transition states in the mammary gland. (A) Mammary gland sections from healthy human patients were stained for the indicated proteins (inset: 3X, n = 4), E-cadherin expression and the percentage of ciliated cells were quantified. Representative results from 3 independent experiments are shown. (B-D) Epithelial-like HMLE cells were treated with 1 μg/mL of doxycycline to induce Snail or Zeb1 expression over 12 days. (B) The morphology of the cells was examined by brightfield microscopy, their expression of the indicated proteins was analyzed by western blot and protein levels were quantified (n = 3, mean ± SEM), (D) their ability to form primary cilia was determined and the percentage of ciliated cells was quantified at the indicated time points (n = 3, mean ± SEM). Representative results from 3 independent experiments are shown. All scale bars: Brightfield 50 μm, Immunofluorescence 15 μm. * p ≤ 0.05, ** p ≤ 0.01, *** p ≤ 0.001.

To determine whether primary ciliogenesis is a transient or a stable response to EMT activation, we used HMLE cells, which are immortalized human mammary epithelial cells (harvested from a mammoplasty and expressing hTERT and SV40 Large T proteins) (*13*). We previously generated HMLE variants in which doxycycline induces ectopic expression of the Snail or Zeb1 EMT-TFs (*11*). To eliminate pre-existing mesenchymal cells from the un-induced HMLE variant populations, we used fluorescence-activated cell sorting (FACS) to isolate the most epithelial cells based on their CD44^Lo^;CD24^Hi^ phenotype (Fig. S1B) (*2*). We then tested our ability to induce EMT and detect various EMT transition states in these epithelial-enriched populations by culturing them with doxycycline for 12 days and examining their morphology and expression of epithelial (E-cadherin) and mesenchymal (Vimentin) markers at various time points. At day 0, both HMLE variants displayed the classic epithelial-like (E-like) cobblestone morphology (Fig. 1B), and lacked detectable expression of either Snail or Zeb1 (Fig. 1C). Doxycycline treatment induced Snail and Zeb1 expression with similar kinetics, both appearing by day 4 (Fig. 1C). Despite these similarities, the two variant populations showed differences in their EMT transitions; the Snail-expressing cells adopted an elongated mesenchymal-like (M-like) morphology progressively between days 8-12, while the Zeb1-expressing cells gained an M-like appearance as early as day 4 and became more distinctly mesenchymal at later time points (Fig. 1B). These changes were concurrent with E-cadherin loss, which was only partial at day 4, and more pronounced at later timepoints in Snail-expressing cells, but was already largely lost 4 days after Zeb1 induction (Fig. 1C). The mesenchymal marker, Vimentin, increased gradually over 12 days in both HMLE variants (Fig. 1C).

Having defined the time windows in which the EMT transition states occurred, we next assessed the representation of primary cilia across these states by staining cells for Arl13b alongside E-cadherin. We observed a significant increase in primary ciliogenesis in response to Snail (day 8) or Zeb1 (starting at day 2 and increasing dramatically on day 4) expression (Fig. 1D), which coincided with the marked reduction of E-cadherin, intermediate expression levels of Vimentin, and a partial M-like phenotype (Fig. 1B-D). Together, these data show that EMT programs induce primary ciliogenesis at intermediate epithelial-mesenchymal transition states.

### EMT-TFs activate the expression of positive regulators of cilium assembly to promote primary ciliogenesis

The mechanisms by which EMT programs induce primary ciliogenesis have been elusive. To determine the underlying molecular mechanisms, we conducted comparative analyses of control (CTL) and Snail-expressing HMLE cells under ciliogenesis-permissive conditions. To do so, CTL and Snail cells were grown to high confluence and serum starved. Replicate samples were subjected to immunofluorescent staining (Fig. S2A) or western blotting (Fig. S2B), which confirmed that the Snail-expressing cells were highly ciliated relative to control cells (Fig. S2A), and expressed lower levels of E-cadherin and higher levels of Fibronectin, N-cadherin, Vimentin, and the Zeb1 and Twist1 EMT-TFs (Fig. S2B). Parallel replicate samples were subjected to RNA-sequencing. This revealed substantial differences in gene expression in CTL versus Snail-expressing cells (Fig. 2A). To identify ciliogenesis regulators whose expression was induced upon EMT induction by Snail, we compared the list of upregulated genes in Snail-expressing versus CTL cells to a cilium gene set, consisting of genes encoding centrosomal and/or ciliary proteins (combined and curated MSigDB GO_Cilium, GO_Cilium Morphogenesis gene sets). We identified 47 genes that overlapped between the two gene signatures, which then represented candidate ciliogenesis inducers and/or ciliary signaling regulators downstream of Snail (Fig. 2B, C).

**Fig. 2:**
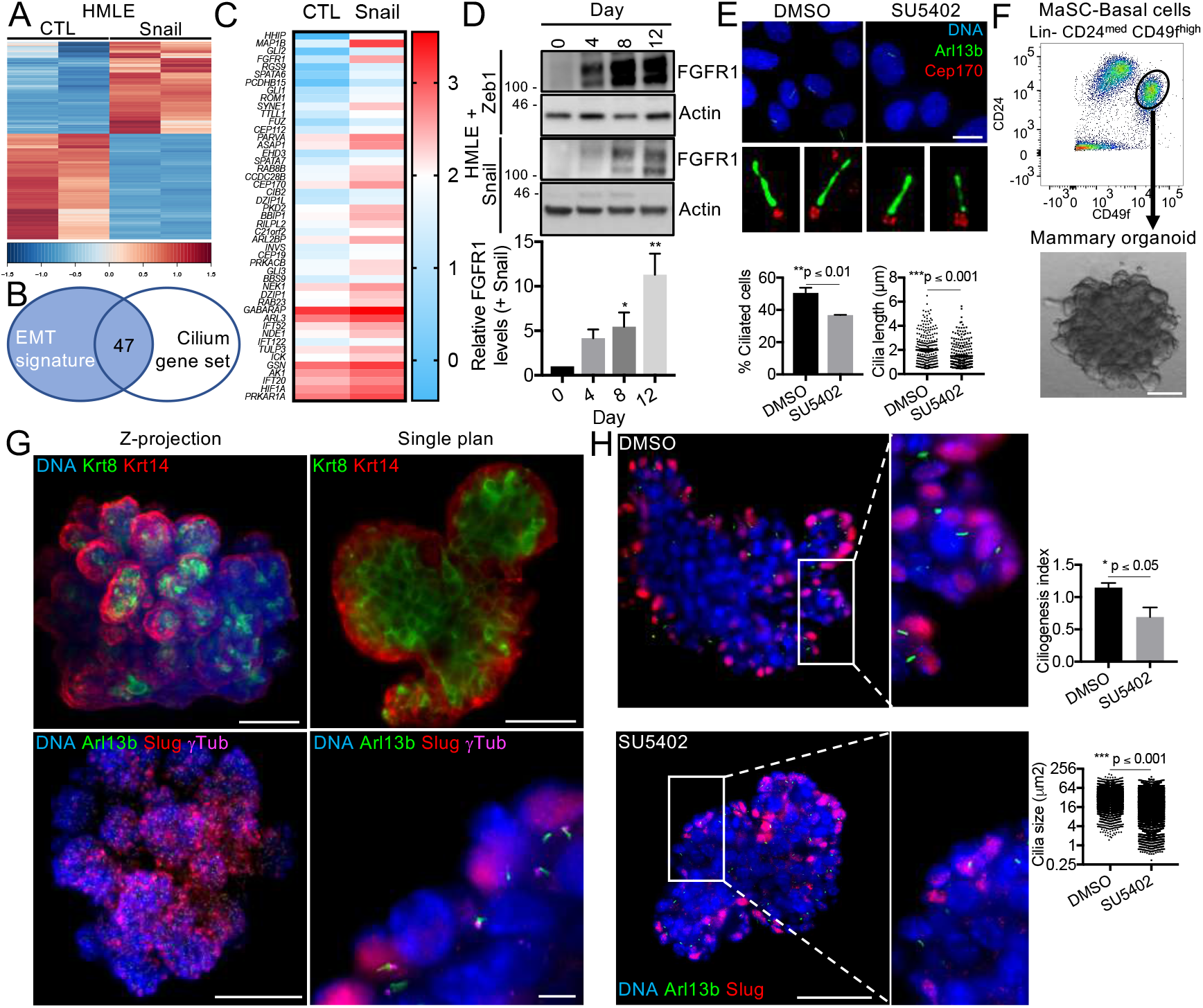
EMT-TFs activate the expression of positive regulators of cilium assembly to promote primary ciliogenesis. (A) Control (CTL) and Snail-expressing (Snail) HMLE cells were grown until high-confluence and serum starved for 24h prior RNA extraction. Gene expression was analyzed by RNA-sequencing. Heat-maps showing the differential expression (DE) genes (q-value ≤ 0.05, Fold-change ≥ 2) between samples are displayed. (B) Venn diagram displaying the overlap between the EMT signature (upregulated genes in Snail-expressing cells relative to CTL cells defined in A) and a curated cilium gene set from MSigDB. 47 genes were found in both gene sets. (C) Heat-map showing the differential expression of the 47 genes between CTL and Snail-expressing HMLE cells. Epithelial-like HMLE cells were treated with doxycycline to induce Snail and Zeb1 expression as described in the legend of Fig. 1 and FGFR1 protein levels were analyzed by western blot and quantified (n = 3, mean ± SEM). Representative results from 3 independent experiments are shown. (E) Snail-expressing HMLE cells were grown until high-confluence and serum starved in the presence of DMSO or of the FGFR1 inhibitor SU5402 (10 μM) for 24h. Treated cells were stained for the indicated proteins and the percentage of ciliated cells and cilia length was measured (n = 3, mean ± SEM). Representative results from 3 independent experiments are shown (inset: 10X, Scale bar: 15 μm). (F-H) MaSC-enriched basal cells (Lin^-^ CD24^med^ CD49f^high^) from C57BL/6J adult females (≥ 10 weeks old) were isolated by FACS using the indicated cell surface markers and plated in 3D. (F) Morphology of mammary organoids was examined by brightfield microscopy. (G) Organoids were stained for the indicated proteins and analyzed by light sheet microscopy. Representative images are shown. Scale bars: upper and lower left panels 100 μm, upper right panel 50 μm and lower right panel 5 μm. (H) Organoids were treated with DMSO or with SU5402 (10 μM), stained for the indicated proteins, and analyzed by light sheet microscopy. Ciliogenesis index (Arl13b particles/Slug particles) as well as cilia size were determined for all conditions (n = 3, mean ± SEM). Representative results from 3 independent experiments are shown. Representative images are shown. Scale bar: 50 μm (inset: 2.5X).

We reasoned that this gene set likely included both direct and indirect Snail targets. To identify potential direct Snail target genes, we reanalyzed the data from an existing Snail ChIP-seq study in mouse mammary tumor cells (*4*), and compared the resulting gene list to our candidate ciliogenesis inducers and cilia components. Remarkably, among these 47 genes, 6 of the 10 most upregulated genes were identified as direct Snail targets: *Fgfr1, Syne1, Rom1, Spata6, Hhip and Map1b* (Fig. S2C, Supplementary Table 1). Importantly, for all 6 of these genes, the peak scores (which represent a normalized value of the read counts identified per gene) were similar to those of known Snail targets (Fig. S2C). The *Fgfr1* gene was particularly interesting. Multiple Snail binding motifs associated with Snail-dependent transcriptional regulation were identified in close proximity to or within the *FGFR1* promoter (Fig. S2D), consistent with the notion that *FGFR1* is directly activated by Snail.

Over the last decade, FGFR1 has emerged as a key inducer of ciliogenesis in the embryonic tissues of lower organisms, including the zebrafish, xenopus, and chick (*14, 15*), and in human lung carcinoma and rhabdomyosarcoma cells (*16*). FGFR1 directly controls intraflagellar transport (IFT) by inducing expression of IFT system components and enabling their import into the cilium (*14, 15*). FGFR1 has not been previously linked to ciliogenesis in the mammary gland, but it is known to cooperate with FGFR2 to promote mammary gland development by regulating stemness of MaSCs (*17*).

Given these observations, we hypothesized that EMT programs activate FGFR1 expression to induce ciliogenesis. After validating the upregulation of *FGFR1* in Snail-expressing versus CTL HMLE cells by real-time qPCR analysis (Fig. S2E), we examined FGFR1 protein expression in Snail- and Zeb1-expressing cells at various EMT transition states using our doxycycline induction scheme. We found that FGFR1 expression was markedly induced by Snail and Zeb1 expression at day 8 and day 4, respectively (Fig. 2D), coinciding with the induction of primary ciliogenesis described earlier (Fig. 1D).

To directly test the role of FGFR1 in mammary ciliogenesis, we cultured our Snail-expressing HMLE cells in ciliogenesis-permissive conditions in the presence of various concentrations of the small-molecule FGFR tyrosine kinase inhibitor, SU5402, or DMSO vehicle control. Initially, we examined the impact of drug treatment on FGFR1 activity by assessing the phosphorylation status of two known FGFR1 downstream targets, AKT and MEK. We saw a significant decrease in the phosphorylation of both targets when cells were treated at 10 μM of SU5402 (Fig. S2F). Importantly, this drug treatment did not alter cell cycle withdrawal (as judged by FACS quantification of G2/M cells; Fig. S2G) or the E-M status of the cells (as judged by western blotting for E-cadherin and Vimentin; Fig. S2H), which could impact ciliogenesis indirectly. We then asked whether it alters ciliogenesis by staining DMSO- and SU5402-treated cells for Arl13b and Cep170 (a centrosome marker). This revealed that FGFR1 inhibition is associated with a significant decrease in the percentage of cells displaying primary cilia and in the length of the cilia that succeeded in forming (Fig. 2E).

To evaluate the role of FGFR1 in ciliogenesis in a more physiological setting, we examined the impact of SU5402 treatment on ciliogenesis in mammary organoids. For this, we isolated MaSC-enriched basal cells from adult female mice by FACS based on their Lin^-^;CD24^med^;CD49f^Hi^ phenotype (Fig. 2F). When plated in appropriate 3D culture conditions, these MaSCs form solid organoids (Fig. 2F) that are mostly clonal, as judged by analyses of mixtures of MaSCs with or without dTomato-expression (Fig. S2I). Importantly, these organoids form rudimentary branches comprised of Krt8^+^;Krt14^-^ cells surrounded by Krt14^+^;Krt8^-^ cells, mimicking the complex cellular architecture of the mammary gland (Fig. 2G). Moreover, the Krt14^+^ cells of the outer layer expressed Slug and displayed primary cilia (Fig. 2G), similar to the human and mouse mammary glands *in vivo* (Fig. 1A, S1A).

We tested the impact of SU5402 in this 3D assay at various concentrations. Consistent with published data(*18*), we found that SU5402 suppresses organoid formation in a dose-dependent manner (Fig. S2J), confirming the importance of FGF signaling for stemness. Additionally, we found that SU5402 treatment significantly reduced the fraction of ciliated cells, as well as cilium length (Fig. 2H). Together, these data showed that EMT-TFs activate the expression of ciliogenesis inducers, including *FGFR1*, which positively regulate primary ciliogenesis in MaSC-enriched basal cells of the mammary epithelium, and that FGF signaling is critical for MaSC stemness.

### Primary cilia control a BBS11-GLIS2 signaling axis

Primary cilia act as cell signaling centers (*19*). To identify signaling pathways that are controlled by primary cilia in mammary epithelial cells, we further expanded our gene expression analysis to Snail-expressing cells grown to high confluence and serum starved in the presence of DMSO or with Ciliobrevin A, a small-molecule ciliogenesis inhibitor (CilA) (*20*). As above, replicate samples of cells were stained for Arl13b and γTubulin to assess the presence of primary cilia under the two conditions (Fig. 3A), confirming that ciliogenesis was significantly inhibited by CilA-treatment relative to DMSO treatment. Replicate samples were then subjected to RNA-sequencing (Fig. 3B). This revealed substantial changes in the gene expression profiles of Snail-expressing cells treated with CilA relative to DMSO-treated cells (Fig. 3B).

**Fig. 3:**
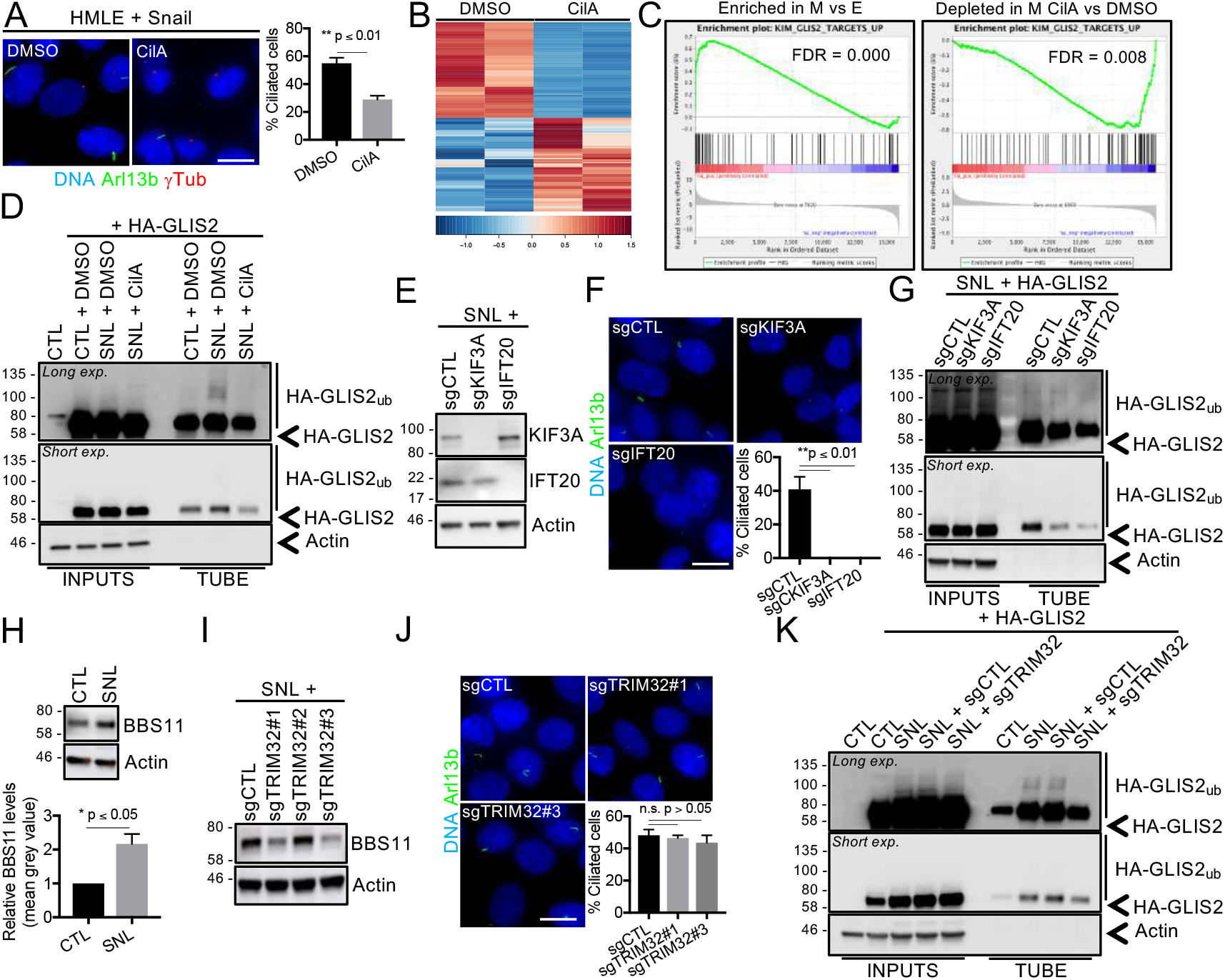
Primary cilia control a BBS11-GLIS2 signaling axis. (A, B) CTL and Snail-expressing HMLE cells were grown until high confluence and serum starved for 24h with DMSO and Ciliobrevin A (CilA). (A) Cells were stained for the indicated proteins and the percentage of ciliated cells was determined. Scale bar: 15 μm (n = 3 mean ± SEM). Representative results from 3 independent experiments are shown. (B) Gene expression was analyzed by RNA-sequencing in both conditions. Heat-maps showing the differentially expressed genes (q-value ≤ 0.05, Fold-change ≥ 2) between samples are displayed. (C) A GSEA analysis shows significant enrichment of GLIS2 target genes in the list of upregulated genes in Snail-expressing cells (M) relative to CTL (E) cells (left panel) and depletion in the list of Snail-expressing (M) CilA-treated relative to DMSO-treated cells (right panel) (False discovery rate (FDR) ≤ 0.05). (D) CTL and Snail-expressing (SNL) HMLE cells that express HA-GLIS2 were treated as described above. Ubiquitinated proteins were purified by TUBE pull-down and analyzed by western blot. Representative results from 3 independent experiments are shown. (E) *KIF3A* and *IFT20* knockouts were validated by western blot of extracts from the indicated cells. (F) The impact on primary ciliogenesis was assessed as described in A. Scale bar: 15 μm. (G) The impact on HA-GLIS2 ubiquitination was analyzed as described in D. (H) BBS11 expression levels were determined in the indicated cells by western blot analysis and quantified (n = 3 mean ± SEM). Representative results from 3 independent experiments are shown. (I) *TRIM32* knockout was validated by western blot of extracts from the indicated cells. (J) The impact on primary ciliogenesis was assessed as described in A. Scale bar: 15 μm. (K) The impact of TRIM32 knockout on HA-GLIS2 ubiquitination was analyzed, as described in D, in the indicated cells.

To identify signaling pathways induced by EMT and primary cilia, we performed gene set enrichment analysis (GSEA). Here, we looked for significantly enriched gene sets that include genes that are upregulated in Snail-expressing HMLE cells versus CTL cells (Fig. 2A) and are downregulated in Snail-expressing CilA-treated cells versus DMSO-treated cells (Fig. 3B). This revealed a gene set that is known to be regulated by GLIS2, a transcriptional repressor (Fig. 3C). As we found, GLIS2 target genes are upregulated in the Snail-expressing cells and repressed upon CilA treatment, suggesting that GLIS2 is inactivated by cilia.

GLIS2 is a transcriptional repressor that is essential for normal development of both the kidney and incisor tooth epithelium (*21-23*). Recent studies showed that it acts as a central regulator of stem and progenitor cell maintenance and differentiation(*23, 24*). Importantly, existing data already suggested a link between GLIS2 and primary cilia. First, GLIS2 loss causes nephronophthisis, a renal ciliopathy (*21*). Second, in the kidney and incisor tooth epithelial cells, GLIS2 localizes to primary cilia (*21, 23*). However, the molecular basis of the interplay between GLIS2 and primary cilia is unknown. Moreover, GLIS2 has not been previously implicated in mammary gland biology. Our own data clearly suggest that primary cilia mediate GLIS2 inhibition in the mammary epithelium.

To explore the underlying mechanism of this repression, we first analyzed *GLIS2* RNA expression in CTL, Snail, Snail+DMSO and Snail+CilA HMLE variants using our RNA-sequencing dataset, as well as by real-time qPCR analysis. Neither the RNA-sequencing analysis nor the RT-qPCR analysis indicated that *GLIS2* expression is repressed upon EMT and ciliogenesis (Fig. S3A, B). Notably, two of the few known GLIS2 target genes, *GLI1* and *CDH11*, were indeed upregulated in Snail versus CTL cells, and downregulated in Snail+CilA versus Snail+DMSO cells (Fig. S3B). Thus, we concluded that GLIS2 is inhibited post-transcriptionally in a cilium-dependent manner.

GLIS2 can undergo ubiquitination-dependent inactivation, a process that is known to require BBS11, an E3-ubiquitin ligase, and a member of the BBS family (*25*). This was particularly intriguing, as BBS stands for Bardet Biedl Syndrome, a ciliopathy (*26*). We therefore asked whether primary cilia inhibit GLIS2 by promoting its ubiquitination. To do so, an HA-tagged version of GLIS2 was introduced in CTL or Snail-expressing HMLE cells, which were then treated with DMSO or CilA. Total proteins were extracted, and ubiquitinated proteins were isolated by affinity chromatography using Tandem Ubiquitin-Binding Entities (TUBE) and then analyzed by western blot.

Comparable levels of total ubiquitinated proteins were recovered in all samples (Fig. S3C). However, we detected a dramatic increase in the levels of ubiquitinated HA-GLIS2 in the Snail+DMSO cells compared to the CTL + DMSO cells (Fig. 3D). Specifically, the lowest mobility band, which we presume is the mono-ubiquitinated form, was present at higher levels in Snail+DMSO versus CTL+DMSO cells (Fig. 3D, short exposure). We also detected multiple, high molecular weight ubiquitinated HA-GLIS2 species that appeared to be unique to the Snail+DMSO cells (Fig. 3D, long exposure). Remarkably, the levels and pattern of ubiquitinated HA-GLIS2 species in the Snail+CilA cells closely resembled that of the CTL+DMSO cells, not Snail+DMSO (Fig. 3D), strongly suggesting that these are cilium-dependent events.

To more directly test whether primary cilia are indeed required for GLIS2 ubiquitination, we examined the ubiquitination status of HA-GLIS2 in Snail-expressing cells in which *KIF3A* and *IFT20*, two essential ciliogenesis regulators, were knocked-out (KO) using CRISPR. To ensure complete gene KO, we selected 5 CRISPR single cell clones with total protein loss and then pooled these knockout clones. Control pools of single cell clones were also generated. We observed that ciliogenesis was abolished in KO cells relative to wild-type control cells (Fig. 3E, F). Genetic inhibition of primary ciliogenesis decreased the levels and pattern of ubiquitinated HA-GLIS2 species (Fig. 3G), in a comparable manner to the pharmacological inhibition of ciliogenesis (Fig. 3D). Together, these data showed that primary cilia induce GLIS2 ubiquitination in mammary epithelial cells.

We then asked if the E3 ubiquitin ligase BBS11 contributes to the observed GLIS2 ubiquitination. We found that BBS11 is significantly upregulated in Snail-expressing HMLE cells, relative to CTL cells, at the level of protein (Fig. 3H) but not mRNA (Fig. S3D, E). We then knocked out the BBS11 gene (*TRIM32*) in Snail-expressing cells using CRISPR (Fig. 3I). Here, two out of the three sgRNAs against *TRIM32*, #1 and #3, yielded cell populations exhibiting a marked reduction of BBS11 protein levels (Fig. 3I). Weak BBS11 protein was still detectable in these populations due to the cell-to-cell variability in the inactivation of TRIM32 in this polyclonal cell population. Next, we grew the sgCTL and the two sgTRIM32 KO populations, #1 and #3, in ciliogenesis-permissive conditions and found that BBS11 loss had no detectable impact on either the E-M status of the cells (Fig. S3 F-G) or on primary ciliogenesis (Fig. 3J). We then examined ubiquitination of HA-GLIS2 in the sgCTL and sgTRIM32#3 KO cells, as described above. In a comparable manner to the pharmacological inhibition or genetic ablation of primary cilia, BBS11 loss reduced the levels and pattern of HA-GLIS2 ubiquitination (Fig. 3K). Hence, we concluded that BBS11 is required for primary cilium-dependent GLIS2 ubiquitination and acts downstream of primary cilium formation.

### GLIS2 is a central regulator of basal MaSC and mammary gland development

Our previously reported data established that EMT-induced primary ciliogenesis in MaSC-enriched basal cells promotes their stemness (*11*). Given this finding, we undertook to test the role of the primary cilium-GLIS2 signaling axis in this process. We initiated this study in our HMLE model, in which the ability to form mammospheres, which are 3D floating structures comprised of undifferentiated cells, serves as a measure of their SC-like properties (*2*).

To begin, we assessed primary cilia representation in mammospheres arising from HMLE cells that have gained the ability to form these structures upon Snail expression, doing so by staining for Arl13b and γTubulin (Fig. 4A, B). This showed that the undifferentiated cells in mammospheres formed primary cilia (Fig. 4B). To investigate the role of GLIS2 in this process, we took advantage of a previously described GLIS2 truncation mutant (*GLIS2* c. 1_444del), termed GLIS2Cter, that can act as a constitutive transcriptional repressor (*27*). Thus, we introduced GLIS2Cter or a control vector into our Snail-expressing HMLE cells (Fig. S4A). We first confirmed the ability of GLIS2Cter to mediate transcriptional repression using qRT-PCR to gauge the expression of the known GLIS2 target genes, *GLI1* and *CDH11*, and found that both transcripts were downregulated, relative to the control vector cells (Fig. S4B). We then showed that GLIS2Cter did not alter the M-like status of these cells, as judged by morphology and expression of E-M markers (Fig. S4A, C), their ability to form primary cilia (Fig. S4D), and their 2D proliferation rate (Fig. S4E). In subsequent mammosphere assays, we found that GLIS2Cter expression significantly inhibited the mammosphere-forming capacity of the Snail-expressing HMLE cells relative to control vector cells (Fig. 4C).

**Fig. 4:**
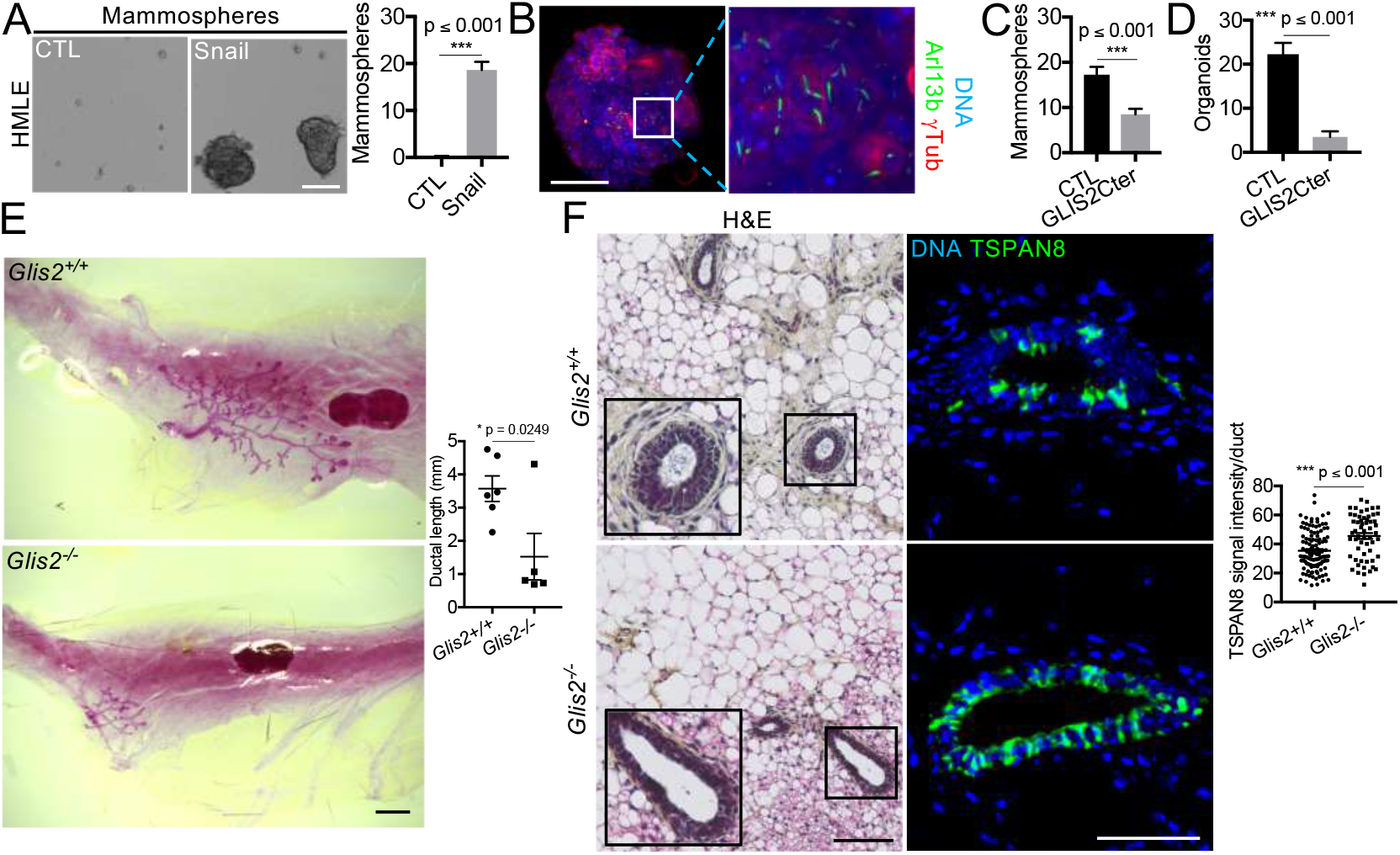
GLIS2 is a central regulator of basal MaSC and mammary gland development. (A-C) Mammospheres from Snail-expressing HMLE cells were examined for morphology by brightfield microscopy or for ciliated cells by immunofluorescence for the indicated proteins by light sheet microscopy. Mammosphere-forming capacity was quantified for the indicated cells (n = 3 mean ± SEM). Representative results from 3 independent experiments are shown. Scale bars: 100 μm. (D) Organoid-forming capacity was determined for sorted MaSC-enriched basal cells in which GLIS2Cter was overexpressed (n = 3 mean ± SEM). Representative results from 3 independent experiments are shown. (E) Mammary glands from *Glis2*^+/+^ or *Glis2*^-/-^ female mice (4 weeks old, n ≥ 5) were stained with Carmine Alum and ductal length was analyzed after whole mount preparation. Scale bar: 1 mm. (F) Paraffin sections from the mammary glands were stained with H&E or for the indicated protein. TSPAN8 signal intensity was measured per duct in all ducts of the sections analyzed. Representative images are shown. Scale bars: H&E 100 μm (inset: 2X), immunofluorescence 50 μm.

Given these findings, we extended our analyses to primary basal MaSCs. Thus, we isolated mouse basal MaSCs by FACS, as described above, and generated populations with ectopic expression of either GLIS2Cter or a control vector. We then examined the ability of these cells to generate organoids and observed a significant reduction in response to GLIS2Cter expression (Fig. 4D). Taken together, these data showed that perturbation of the primary cilium-GLIS2 axis results in inhibition of stem cell-like cells in both the HMLE population and the basal murine MaSCs and thus impairs *ex vivo* organogenesis.

We reasoned that if Glis2 were important for regulation of MaSCs *in vivo*, its deletion should perturb mammary gland development in mice suffering from germline KO of the *Glis2* gene. We further hypothesized that the resulting mammary defects would reflect inappropriate expansion of MaSCs, at the expense of cell differentiation and proper lineage commitment. To test these predictions, we examined mammary gland development in 4-week-old *Glis2*^+/+^ and *Glis2*^-/-^ female mice, through carmine-alum staining of whole-mount mammary glands. In agreement with our hypothesis, ductal morphogenesis was profoundly impaired in *Glis2*-deficient mice, as evidenced by the presence of only small mammary rudiments relative to the *Glis2*^+/+^ controls (Fig. 4E). Additionally, H&E staining of paraffin sections showed that *Glis2*^+/+^ glands had the expected bilayered epithelium, but this organization was disrupted in *Glis2*^-/-^ mice (Fig. 4F). Most importantly, *Glis2*^-/-^ rudimentary ducts were enriched for TSPAN8-expressing cells relative to *Glis2*^+/+^ controls (Fig. 4F). TSPAN8 is a plasma membrane protein that is highly expressed in basal MaSCs and to a lesser extent in luminal progenitor cells of the mammary epithelium (*28*). Collectively, our data showed that GLIS2 regulates the expansion of basal MaSCs in a manner that is required for normal mammary gland development.

### Primary cilium-dependent regulation of GLIS2 controls claudin-low MaTIC stemness and tumorigenicity

We have previously shown that the role of primary cilia is not only important for normal basal MaSCs but also promotes the tumor-initiating capacity of MaTICs in an orthotopic murine carcinoma model (*11*). This analysis was conducted using HMLE cells transformed with H-RASG12V (called HMLER), which, upon E-cadherin knockdown, become more M-like (Fig. S5A) and acquire MaTIC properties (*11*). These cells can form tumorspheres comprised of non-differentiated cells that display cilia *in vitro* (Fig. 5A-B) and generate ciliated tumors that display the hallmarks of claudin-low breast cancers (Fig. S5B).

**Fig. 5:**
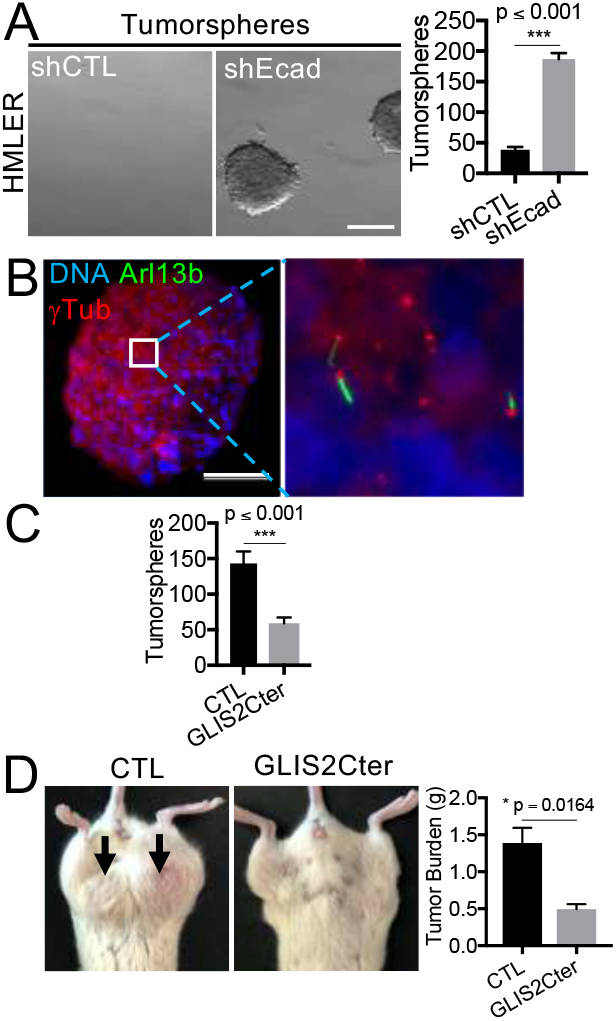
The primary cilium-GLIS2 axis controls claudin-low MaTIC stemness and tumorigenicity. (A-C) Tumorspheres from shEcad-expressing HMLER cells were examined for morphology by brightfield microscopy or for ciliated cells by immunofluorescence for the indicated proteins and light sheet microscopy. Tumorsphere-forming capacity was quantified for the indicated cell variants (n = 3 mean ± SEM). Representative results from 3 independent experiments are shown. Scale bars: Brightfield image, 300 μm; and immunofluorescence image, 100 μm. (D) Bilateral orthotopic implantations were conducted with HMLER shEcad control (CTL) or GLIS2Cter-expressing cells. Representative mice are shown. Tumor burden per mouse (mean ± SEM) was determined 8 weeks post-implantation with two sites of implantation per mouse and four mice per cell type.

We therefore asked whether primary cilia control GLIS2 in these transformed HMLER cells. We began by assessing the cilium-dependent gene expression events in HMLER variants. For this, we grew HMLER shCTL and shE-cadherin (shEcad)-expressing cells in ciliogenesis-permissive conditions, doing so with or without DMSO and CilA. For each condition, samples were stained for Arl13b and E-cadherin to assess primary cilia representation. These analyses confirmed that E-cadherin loss in HMLER cells led to a dramatic increase in the percentage of ciliated cells, relative to shCTL cells (Fig. S5C), and that this ciliogenesis was reduced significantly by treatment with CilA (Fig. S5C). Parallel samples were analyzed by RNA-sequencing, revealing substantial changes in gene expression profiles between these HMLER populations (Fig. S5D). Importantly, GSEA revealed that the GLIS2 target genes were upregulated in shEcad-expressing vs shCTL cells and downregulated in shEcad-expressing CilA vs DMSO treated cells (Fig. S5E). Thus, GLIS2 is inhibited in a cilium-dependent manner in transformed HMLER cells enriched for MaTICs in a similar manner to that observed in their non-transformed HMLE counterparts.

We next asked whether GLIS2 activity influences the properties of these HMLER cells, doing so by generating HMLER shEcad populations expressing GLIS2Cter. As with Snail-expressing HMLE cells, this perturbation did not alter the E-M status, primary ciliogenesis, or proliferation rate of the HMLER+shEcad cells in monolayer culture (Fig. S5F-I). However, GLIS2Cter did repress *GLI1* and *CDH11* gene expression (Fig. S5J), validating its activity. Importantly, GLIS2Cter expression decreased the tumorsphere-forming capacity of HMLER shEcad cells relative to the control vector cells (Fig. 5C).

To further validate the key role of GLIS2 in tumors, we also compared the tumorigenic capacity of GLIS2Cter-expressing and CTL HMLER shEcad cells in orthotopic tumor implantation experiments. Consistent with our *in vitro* assays, the GLIS2Cter expression significantly reduced the *in vivo* tumorigenic capacity of the HMLER shEcad CTL cells (Fig. 5D). These data show that the primary cilium-dependent regulation of GLIS2 is required for the proliferative and tumorigenic capacities of MaTICs, which form tumors displaying hallmarks of claudin-low breast cancers.

To further analyze primary ciliogenesis in claudin-low human breast cancers, we established a tumor biobank by characterizing tumor samples in a previously assembled TNBC biobank. We extracted total RNA from sections of frozen tumors (n = 62) and analyzed gene expression by RNA-sequencing. Among the 62 TNBCs, 23 claudin-low tumors were then identified based on their differential expression of a panel of genes (Fig. 6A), previously identified by Prat and colleagues (*29*). We were able to stain paraffin sections for 20 of them, along with 3 luminal B tumors (as controls) for H&E and for E-cadherin, Claudin 4, and Claudin 7.

**Fig. 6:**
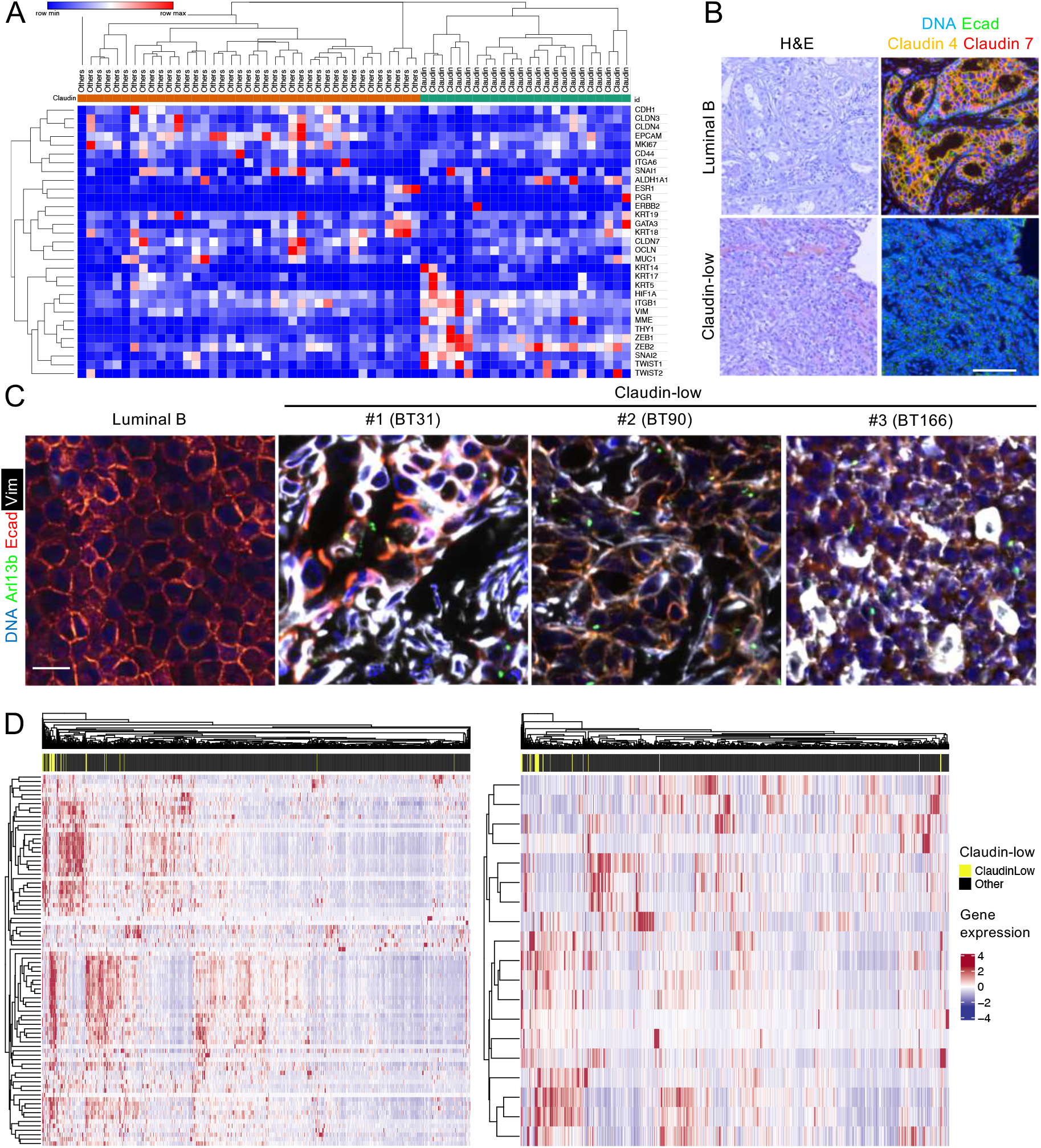
Primary cilia mark claudin-low tumor cells in E-M states and high expression of GLIS2 target genes is a hallmark of claudin-low tumors. (A) Human breast tumor fragments from 62 independent patients were flash frozen after surgery. RNA was extracted from sections of frozen tumors and gene expression was analyzed by RNA-sequencing. Expression of a subset of genes was analyzed to discriminate claudin-low tumors from other molecular subtypes. The Heat-map showing the differential expression of the genes is displayed. (B-C) Paraffin sections of breast tumors were stained with H&E or for the indicated proteins. Scale bar: 100 μm (B), 20 μm (C). (D) Heat maps showing the differential expression of GLIS2 target genes (left panel), and the reference list of genes used to mark claudin-low tumors (right panel), in breast cancers are displayed, along with their ability to segregate claudin-low tumors from other breast tumors.

We found that, in contrast to the luminal B tumors, which all demonstrated the typical appearance of differentiated breast cancers, the claudin-low tumors displayed a poorly differentiated phenotype (Fig. 6B). Luminal tumors expressed higher levels of epithelial markers than did 85% of the claudin-low tumors (Fig. 6B, S6A). We also co-stained luminal and claudin-low tumor sections for Vimentin, E-cadherin, Arl13b, and γTubulin in order to assess primary cilia representation. This revealed ciliated cancer cells in claudin-low tumors that co-expressed E-cadherin and Vimentin, but not in luminal tumors, indicating that they reside in an intermediate EMT transition state (Fig. 6C, S6B). Moreover, these cells represented a small fraction of overall tumor cells, consistent with the hypothesis that the ciliated claudin-low tumor cells are MaTICs.

To determine whether high expression of the GLIS2 signature is a hallmark of claudin-low tumors across breast cancers in general, we compared the expression of GLIS2 target genes in 1082 breast cancers, using a public dataset from The Cancer Genome Atlas (TCGA) (*30*). Remarkably, we found that high expression of GLIS2 target genes enables the segregation of claudin-low tumors from other breast tumors in a manner similar to the Prat et al reference panel of genes used to mark claudin-low cancers (Fig. 6D) (*29*). Moreover, GSEA revealed that the GLIS2 gene set is significantly enriched in the signature that was initially used to define the claudin-low tumors (Fig. S6C) (*29*). Collectively, our data showed that primary cilia are assembled in claudin-low breast tumor cells that are enriched in MaTICs, in which they control GLIS2 inactivation to promote the tumorigenic capacity of these cells. Furthermore, high expression of GLIS2 target genes serves as a hallmark of claudin-low breast cancers.

## Discussion

Our previously reported data established that EMT programs induce ciliogenesis in mammary epithelial cells (*11*). However, we lacked a mechanistic understanding of the link between EMT and primary ciliogenesis, including the degree to which ciliogenesis is a stable or a transient response upon EMT activation, as well as the underlying molecular mechanisms. Our data reinforce the connection between EMT programs and primary ciliogenesis both in the human and mouse mammary glands. Specifically, we show that the propensity to form primary cilia closely follows entrance into mixed epithelial-mesenchymal phenotypic states and that primary cilia are assembled in mammary epithelial cells that display an array of M-like phenotypes. While we do not see a decrease in the percentage of ciliated cells in the most mesenchymal phenotypic states in our systems, we do not exclude the possibility that primary ciliogenesis is repressed at more extreme mesenchymal EMT transition states. Indeed, this latter hypothesis is supported by an elegant study of Epithelial-Myofibroblast Transition (EMyT), a more extreme type of EMT that can occur in kidney epithelial cells (*31*). Here, primary cilia are induced during early transition states, where they promote EMyT progression but are ultimately repressed when the cells acquire the highly mesenchymal, myofibroblast phenotype (*31*).

In yet another context, the murine coronary vasculature, a link between EMT and primary ciliogenesis was recently identified during development (*32*). Here, the authors found that epicardial cells that undergo EMT assemble primary cilia, and that genetic disruption of ciliogenesis in these cells perturbed EMT and migration, causing coronary artery developmental defects (*32*).

These findings cement a strong connection between EMT and primary ciliogenesis in multiple contexts and underscore the concept that EMT is not a simple binary switch between E and M states, but instead generates a spectrum of transitional phenotypic states that can have a distinctive relationship with the primary cilium biology. Our data established that EMT induces primary ciliogenesis at intermediate and pronounced transition states in mammary epithelial cells of the human and mouse mammary gland to support stemness.

The precise molecular mechanisms by which EMT programs induce primary ciliogenesis were previously unknown. Our data show that EMT-TFs activate the transcription of ciliogenesis inducers in mammary epithelial cells. Specifically, we found that FGFR1 expression is induced by EMT-TFs at intermediate epithelial-mesenchymal transition states. We show that FGFR1 is responsible, at least in part, for primary cilium assembly and elongation upon activation of EMT programs. A recent study found that FGFR1 promotes ciliogenesis in the hair cells of the inner ear of the chick embryo by phosphorylating the protocadherin Pcdh15 (*15*). Interestingly, we found another member of the protocadherin family, *PCDHB15*, in our identified list of 47 putative ciliogenesis inducers, together with *FGFR1* (Fig. 2C), raising the intriguing possibility that FGFR1 cooperates with Protocadherin beta-15 in promoting primary ciliogenesis in mammary epithelial cells. Of note, FGFR1 cooperates with FGFR2 to promote mammary gland development and regeneration by enabling the stemness of MaSCs (*17, 18*). Our own observations show that FGFR inhibition represses both ciliogenesis and organoid-forming capacity in MaSCs. Collectively, these data suggest that FGFR1 controls ciliogenesis, thereby enabling basal MaSC stemness and mammary gland development.

Our prior work revealed that primary cilia control Hedgehog signaling in mammary cells (*11*). In the current study, we used gene expression analyses to gain a broader and unbiased insight into the signaling pathways that depend on the primary cilium in mammary cells; these analyses revealed that cilia control a GLIS2 signaling axis. This was highly gratifying because, although GLIS2 is a relatively understudied protein, it had already been linked to EMT, primary cilia, and stemness in other contexts. With regard to EMT, GLIS2 was found by others to repress EMT in the kidney (*21*). We did not observe evidence of increased Mesenchymal-Epithelial Transition (MET) when GLIS2 was ectopically activated in M-like mammary cells, leading us to conclude that GLIS2 acts downstream of EMT in the mammary epithelium. However, our data do not rule out the possibility that GLIS2 inactivation helps reinforce EMT through a positive feedback loop. With regard to primary cilia, GLIS2 was shown by others to localize to the primary cilium in incisor tooth epithelial stem cells undergoing differentiation, and in kidney cells (*21, 23*). However, the role of the primary cilium in regulating GLIS2 had not been previously addressed. Our data show that GLIS2 is ubiquitinated and inhibited in a cilium-dependent and BBS11-dependent fashion. Notably, several members of the BBS family of proteins are known to bind to GLIS2, including BBS11, and these various BBS proteins have been found to localize to primary cilia or centriolar satellites (*25, 26, 33*). Thus, we speculate that primary cilia serve as cell signaling platforms where BBS11 induces GLIS2 ubiquitination and consequent inactivation. Finally, and most importantly, our data show that GLIS2 regulates stemness of basal MaSCs. The signaling pathway(s) downstream of GLIS2 that control(s) MaSC-stemness remain to be investigated, but it is intriguing to note that GLIS2 is known to repress two key stem cell signaling pathways, Wnt and Hedgehog (*34, 35*). This intersects with our prior finding that Hedgehog signaling acts downstream of primary ciliogenesis in basal MaSCs and MaTICs (*11*).

As mentioned, prior reports revealed that GLIS2 controls normal development of both the kidney and the incisor tooth epithelium (*21-23*). Our own data show that GLIS2 also plays a critical role in normal mammary gland development, which is consistent with our discovery of its role in MaSC regulation. More specifically, we found a major morphogenetic defect at puberty in *Glis2*-deficient female mice. We detected only rudimentary trees in *Glis2*-deficient animals, in which the normal bilayer structure of the mammary epithelium is disrupted and an excess of basal MaSCs was detected. These findings suggest that there is an inappropriate persistence or expansion of the SC compartment and/or a failure of engagement of the differentiation program at the appropriate developmental time.

Our prior work revealed a critical role for primary cilia in MaTICs of the more mesenchymal HMLER cell line, which yield tumors that display the hallmarks of claudin-low human breast cancers (*11*). Here, we found that primary cilia control GLIS2 in these MaTICs, thereby enabling their self-renewal and tumorigenic capacities both *in vitro* and *in vivo*. To directly address the relevance of these findings in human breast tumors, we established a biobank of claudin-low breast tumor samples. Subsequently, we found that claudin-low breast tumors are enriched in tumor cells that display an M-like phenotype and assemble primary cilia. Significantly, we found that a GLIS2 signature identifies claudin-low tumors from other breast cancer subtypes as effectively as the previously identified claudin-low signature. In conclusion, by establishing molecular mechanisms by which EMT induces ciliogenesis and thereby controls MaSC and MaTIC biology, our findings reveal a novel aspect of the cell biology critical to the development of the mammary gland and the formation of claudin-low breast cancers.

## Materials and Methods

### Animals

Transgenic and wild-type mice were housed and handled in accordance with protocols approved by the Animal Care Committees of the Massachusetts Institute of Technology (USA), the University of Iowa (USA) and the University of Rennes (France). Slug-IRES-YPF and Glis2 mutant mice were generated and selected as previously described (Guo et al., 2012, Kim et al., 2008). C57BL/6J and NOD SCID animals were obtained from the Jackson laboratory (stock numbers 000664 and 001303).

### Human samples

Patient reduction mammoplasty tissue samples and breast tumor samples were obtained in compliance with all relevant laws. Breast cancer patients (n = 62) were diagnosed and treated at the Centre Eugène Marquis between 2013 and 2018, none received chemotherapy, endocrine therapy, or radiation therapy prior to surgery. Tissues were collected by a pathologist after resection by a surgeon. Some fragments were paraffin-embedded (normal tissues and tumors), other fragments were flash-frozen (tumors), and other fragments were dissociated within 2 h after surgical resection (tumors). Briefly, breast tumor pieces were weighed and cut into small fragments (< 2 mm^3^), which were then dissociated with a tumor dissociation kit (Human) using a mechanical–chemical–mechanical cycle in a gentleMACS Dissociator (Miltenyi Biotec), according to the manufacturer’s recommendations. Next, a cycle of mechanical–chemical–mechanical dissociation was performed and dissociated cells were resuspended in RPMI. Macroscopic pieces were removed using a Corning® cell strainer (70 μm). Tumor cells were then washed twice in RPMI (20 ml) and counted using a hemocytometer. Viable cells were then cryopreserved.

### Primary mouse mammary epithelial cell isolation and FACS

Mammary glands from 10-16 weeks old C57BL/6J females were minced and dissociated using collagenase/hyaluronidase (STEMCELL Technologies) diluted (1:10) in DMEM/F-12 medium at 37°C for 5 h under constant agitation. The dissociated glands were spun down at 450 x g for 5 min and resuspended in ammonium chloride solution (STEMCELL Technologies) diluted (4:1) in HBSS buffer supplemented with 10 mM HEPES and 2 % FBS (HF buffer). The samples were spun down and resuspended in warm Trypsin-EDTA (Stem Cell Technologies) by pipetting for 3 min. Trypsin was inactivated by addition of HF buffer. The digested samples were spun down and further digested with dispase solution (STEMCELL Technologies) supplemented with 0.1 mg/mL DNase I for 1 min. Cell suspensions were diluted with HF buffer and filtered through a 40 μm cell strainer to collect single cells. To separate various cell populations, single cells were stained with a Live-Dead fixable violet dye and antibodies against TER-119, CD31, CD45 (PE/Cy7, Biolegend); CD24 (PE, Biolegend); CD49f (APC, Biolegend). Stained cells were sorted on a FACSAria II and FACS Fusion sorters.

### 2D cell culture and treatments

Primary mammary epithelial cells were cultured in EpiCult-B mouse medium (STEMCELL Technologies) supplemented with proliferation supplements, 10 ng/mL recombinant human epidermal growth factor, 10 ng/mL recombinant human basic fibroblast growth factor, 4 μg/mL heparin, and penicillin/streptomycin. HMLE cells were cultured in a 1:1 mixture of DMEM/F-12 supplemented with 10 % FBS, 0.01 mg/mL insulin, 0.48 μg/mL hydrocortisone and complete MEGM supplemented with bovine pituitary hormone (Lonza). For activating tetracycline-inducible Snail and Zeb1 expression, cells were grown with 1 μg/mL doxycycline hyclate (Sigma-Aldrich) in media. For all ciliogenesis assays, cells were grown until high confluence and DMEM/F12 was used to serum starve the cells. For FGFR1 inhibition, cells were treated with SU5402 (Sigma-Aldrich) in DMEM/F12 medium.

### Mammary organoid culture

Matrigel organoid culture was performed as described previously (Guen et al., 2017). Briefly, freshly isolated mammary epithelial cells or transduced cells were cultured in complete Epicult-B medium (STEMCELL Technologies) containing 5 % matrigel (Corning), 5 % heat-inactivated FBS, 10 ng/mL EGF, 10 ng/mL FGF, 4 μg/mL heparin, 5 μM Y-27632. Cells were seeded at 2000 cells/well in 96-well ultralow attachment plates (Corning). Organoids were counted 7-14 days after seeding. For FGFR1 inhibition, SU5402 was added to the medium at 10 μM.

### Mammosphere and tumorsphere culture

Cells were cultured in complete Mammocult medium (STEMCELL Technologies) containing 4 μg/mL heparin, 0.48 μg/mL hydrocortisone, 1% methyl cellulose. Non-transformed cells were seeded at 1000 cells/well and transformed cells at 150 cells/well in 96-well ultralow attachment plates (Corning). Spheres were counted 7-14 days after seeding.

### Mammary gland whole-mount analysis

Glands were fixed in Carnoy’s fixative (100% EtOH, chloroform, glacial acetic acid; 6:3:1) for 4 hours, washed in 70% ethanol for 15 min and gradually rehydrated in 70%, 35%, 15% ethanol baths for 5 min each, and stained with carmine alum overnight at room temperature. Glands were washed with sequential 5 min washes in 50%, 70%, 95%, and 100% ethanol, cleared in xylene overnight and stored in methyl salicylate until analysis using a Nikon macroscope. Images were processed with ImageJ.

### Tumorigenesis assay

For orthotopic cell implantations, tumor cells were resuspended in a 1:1 mixture of complete MEGM medium with matrigel. Cells in 20 μL were injected bilaterally into inguinal mammary fat pads of 8-weeks-old NOD SCID females. Tumors were resected 8 weeks post-implantation and their mass was determined to establish the tumor burden per animal.

### Immunohistochemistry, Immunofluorescence and image analysis

5 μm paraffin tissue sections of formalin-fixed mammary glands and breast tumors were stained following the RUO DISCOVERY Universal staining procedure (Roche) using a Discovery ULTRA staining module. Cells fixed in 4% paraformaldehyde for 10 min on glass coverslips were permeabilized with 0.3% Triton X100 for 10 min and blocked with 5% goat serum for 1 h before staining 1 h with primary antibodies. Organoids and spheres fixed in 4% paraformaldehyde for 30 min were permeabilized with 0.3% Triton X100 for 30 min and blocked with 5% goat serum for 1 h 30 before staining overnight at 4°C with primary antibodies. The following primary antibodies were used: Arl13b (NeuroMab 73-287, 1:100), YFP (Cell Signaling 2956, 1:100), Slug (Cell Signaling 9585, 1:100), acetylated-tubulin (Cell Signaling 5335, 1:500), γ-tubulin (Sigma T5326, 1:50), SMA (Abcam ab21027, 1:100), E-cadherin (Cell Signaling 3195, 1:200), Cep170 (Fisher Scientific 10383523, 1:200), Krt8 (DSHB TROMA-I-s, 1 :1000), Krt14 (Biolegend 905301, 1:1000), Claudin 4 (Fisher Scientific 10303233, 1:200), Claudin 7 (Fisher Scientific 10537403, 1:200), Large T (Santa Cruz sc-20800, 1:200), VIM (Dako M0725, 1:200), TSPAN8 (Personal gift from Jane Visvader, 1:500). The following secondary antibodies were used: anti-mouse 488 (Life Technologies A11001, 1:500), anti-mouse 546 (Life Technologies A11003, 1:500), anti-mouse IgG1 647 (Life Technologies A21240, 1:500), anti-mouse IgG2A 488 (Life Technologies A21131, 1:500), anti-rabbit 488 (Life Technologies A21206, 1:500), anti-rabbit 546 (Life Technologies A11010, 1:500), anti-rabbit 647 (Life Technologies A31573, 1:500), anti-rat 568 (Life Technologies A11077, 1:500). Mounted coverslips with cells were examined using Deltavision Olympus X71 microscopes. Z-stacks were deconvolved (Softworx) and processed with ImageJ. Organoids and spheres were embedded in low-melting point agarose and analyzed using a Light sheet Z1 Zeiss microscope. Z-stacks were analyzed with Imaris and ImageJ.

### Ubiquitination assay

HMLE variants were washed with PBS prior to protein extraction with lysis buffer (PBS, protease inhibitor cocktail (Sigma-Aldrich), phosphatase inhibitor cocktail (Sigma-Aldrich), 0.1 % NP-40, 100 μM PR-619). The pull-down of ubiquitinated proteins from protein extracts was performed with TUBE 2 agarose beads (LifeSensors), following the manufacturer’s instructions. Briefly, protein extracts were incubated for 30 minutes on uncoupled agarose to remove non-specific binding and 2 hours on equilibrated TUBE 2 agarose on a rotating wheel. Beads were washed 3 times with lysis buffer. Proteins were heat-denatured and eluted in laemmli buffer and analyzed by western blot.

### Western blot experiments and cell cycle analysis

These experiments were conducted using standard procedures as described previously (*36, 37*). Western blots were performed using the primary antibodies against Snail (Cell Signaling 3879, 1:500), Twist (Abcam ab50887, 1:200), Zeb1 (Santa Cruz Biotechnology sc-25388, 1:300), E-cadherin (Cell Signaling 3195, 1:1000), Vimentin (Dako M0725, 1:1000), Fibronectin (BD Biosciences 610077, 1:1000), N-cadherin (BD Biosciences 610920, 1:1000), KIF3A (Proteintech 13930-1-AP, 1:1000), IFT20 (Proteintech 13615-1-AP, 1:100), FGFR1 (Abcam ab76464, 1:1000), BBS11 (Proteintech 10326-1-AP, 1:1000), HA (Roche 11867423001, 1:1000), RFP (Rockland 600-401-379S, 1:1000), AKT (Cell Signaling 9272, 1:1000), pAKT (Cell Signaling 4060, 1:1000), MEK (Cell Signaling 9122, 1:1000), pMEK (Cell Signaling 9121, 1:1000), Ub (P4D1) (Santa cruz sc-8017, 1:300), Actin (Sigma-Aldrich 20-33, 1:1000) and secondary antibodies HRP-coupled anti-mouse or anti-rabbit (GE Healthcare and Jackson Laboratories, 1:5000). The cell cycle analysis was performed using HMLE cells that were fixed in 70% ethanol for 1 h on ice. DNA was stained with Propidium Iodide/RNAse staining buffer (BD Biosciences) for 15 min. DNA content of 30,000 cells for each condition was determined using a BD Fortessa X20 flow cytometer and the FlowJo software.

### Quantitative Real Time-PCR

Total RNA was isolated using the PicoPure RNA Isolation Kit (Thermo Fisher Scientific) for mouse cells or the RNeasy kit (Qiagen) for human cells, and cDNAs were generated with random primers and SuperScript III Reverse Transcriptase (Life Technologies). Real-time quantitative PCR reactions were performed with StepOnePlus Real-Time PCR system (Life Technologies) and 7900HT Fast Real-Time PCR System (Applied Biosystem) using SYBR Green PCR Master Mix (Life Technologies) and gene-specific primers. Relative expression levels were normalized to GAPDH. Primers used for qPCR analysis are the following:

Cdh1_fw CAGTGAAGCGGCATCTAAAGC

Cdh1_rev TTGGATTCAGAGGCAGGGTCG

Zeb1_fw CACAGTGGAGAGAAGCCATAC

Zeb1_rev CACTGAGATGTCTTGAGTCCTG

Slug_fw CTCACCTCGGGAGCATACAG

Slug_rev GACTTACACGCCCCAAGGATG

Snai1_fw CACACTGGTGAGAAGCCATTC

Snai1_rev CTTGTGGAGCAAGGACATGCG

Krt14_fw CAGCAAGACAGAGGAGCTGAACC

Krt14_rev CCAGGGATGCTTTCATGCTGAG

Krt18_fw GGTCTCAGCAGATTGAGGAGAG

Krt18_rev CCAAGTCAATCTCCAAGGTCTGG

FGFR1_fw GGAGGCTACAAGGTCCGTTA

FGFR1_rev TGCAGGTGTAGTTGCCCTTG

CDH1_fw CAGTCAAAAGGCCTCTACGG

CDH1_rev CAGAAACGGAGGCCTGATGG

VIM_fw GGAAATGGCTCGTCACCTTC

VIM_rev GAAATCCTGCTCTCCTCGCC

GLI1_fw CACAAGTGCACGTTTGAAGGG

GLI1_rev CATGTATGGCTTCTCACCCG

CDH11_fw CGCAGAGCGTATACCAGATG

CDH11_rev CCTCCTGTGTTTCATAGTCCG

TRIM32_fw CAGTTAACGTGGAAGATTCC

TRIM32_rev GAGGCACTGCTGGATATTGG.

### Plasmids, lentivirus production, CRISPR mutations

KIF3A, IFT20, TRIM32 guide RNAs were selected from http://crispr.mit.edu/ and https://chopchop.cbu.uib.no/ websites and cloned into lentiCRIPRV2 plasmids containing, either a puromycin-resistance gene, or a blasticidin-resistance gene designed to replace the puromycin one. GLIS2 full-length and C-terminal (c. 1_444del) were amplified from a Precision LentiORF GLIS2 plasmid (Horizon Discovery) and cloned in frame with 3xHA or dTomato tags in a pEF_BSD lentiviral plasmid through GIBSON cloning. The following primers were used:

CRISPR

sgCTL_fw GCTGATCTATCGCGGTCGTC

sgCTL_rev GACGACCGCGATAGATCAGC

sgKIF3A_fw GAAATCAATGTGCTACAAAC

sgKIF3A_rev GTTTGTAGCACATTGATTTC

sgIFT20_fw CCAGCAGACCATAGAGCTGA

sgIFT20_rev TCAGCTCTATGGTCTGCTGG

sgTRIM32#1_fw CACCGGCGGACACCATTGATGCTAC

sgTRIM32#1_rev AAACGTAGCATCAATGGTGTCCGCC

sgTRIM32#2_fw CACCGAACTCGTCTGCGGGAACTTA

sgTRIM32#2_rev AAACTAAGTTCCCGCAGACGAGTTC

sgTRIM32#3_fw CACCGGTCTGCCCCGGCAATTCTGC

sgTRIM32#3_rev AAACGCAGAATTGCCGGGGCAGACC

sgTrim32#1_fw CACCGGCGGACGCCATTGATGCTGC

gTrim32#1_rev AAACGCAGCATCAATGGCGTCCGCC

sgmTrim32#3_fw CACCGGGCTGCCTCGGCAGTTCTGC

sgmTrim32#3_rev AAACGCAGAACTGCCGAGGCAGCCC

HA-GLIS2 cloning:

1_GTGTCGTGAgccaccatggtgTACCCATACGATGTTCCTGAC

2_GTCAGGAACATCGTATGGGTAcaccatggtggcTCACGACAC

3_GACGTTCCAGATTACGCTCACTCCCTGGACGAGCCG

4_CGGCTCGTCCAGGGAGTGAGCGTAATCTGGAACGTC

5_GACGTTCCAGATTACGCTTTCCTTACCCCTCCCAAGGAC

6_GTCCTTGGGAGGGGTAAGGAAAGCGTAATCTGGAACGTC

7_CTCAAACCGGCTGTGGTGAACGGATCCGGCGCAACAAACTTC

8_GAAGTTTGTTGCGCCGGATCCGTTCACCACAGCCGGTTTGAG

GLIS2Cter cloning:

1_ CTCAAACCGGCTGTGGTGAACGGATCCGGCGCAACAAACTTC

2_GAAGTTTGTTGCGCCGGATCCGTTCACCACAGCCGGTTTGAG

3_ctctggttatgtgtgggagggctaagaattcgttccggagtcgtcg

4_cgacgactccggaacgaattcttagccctcccacacataaccagag

5_ggatctggagcaacaaacttcACCTTCCTTACCCCTCCCAAGGAC

6_GTCCTTGGGAGGGGTAAGGAAGGTgaagtttgttgctccagatcc

Lentiviruses were produced in 293FT cells after their transfection with lentiCRISPR or pEF_BSD_HA-GLIS2 or pEF_BSD_dTomGLIS2Cter (transfer) in combination with psPAX2 (packaging) and pMD2.G (envelope) plasmids, using Lipofectamine 2000 (Life Technologies) according to the manufacturer’s instructions. Supernatants containing lentiviruses were collected 48 and 72 h post-transfection. Primary mammary epithelial cells, HMLE and HMLER cells growing in a monolayer were transduced with the supernatants-containing viruses in the presence of 8 μg/mL polybrene. Transduced cells were selected with 2 μg/mL puromycin (GIBCO) or 8 μg/mL blasticidin (GIBCO). HMLE *KIF3A*^*-/-*^ and *IFT20*^*-/-*^ clones were selected by FACS.

### Gene expression and ChIP-sequencing analyses

For gene expression analysis in HMLE and HMLER cells, total RNA was extracted from highly confluent cells. 25 ng for each of the two biological replicates per condition was used to generate libraries for whole-transcriptome analysis following High Throughput 3’ Digital Gene Expression library preparation. Libraries were sequenced on an Illumina Nextseq500. Raw sequence reads were collapsed by sequence identity and UMI (Unique Molecular Identifier) and aligned to the human genome (UCSC hg19 build) using TopHat v. 2.0.4. Genic read counts were quantified using the End Sequence Analysis Toolkit (ESAT v1). Count normalization and differential analyses were conducted using the DESeq package (R/Bioconductor) and genes with q-value < 0.05 and absolute fold change greater than 2 were tagged as differentially expressed. Row-normalized heatmaps for differentially expressed genes were created using Heatplus (R/Bioconductor). Gene set enrichment analyses (GSEA) were performed with the GSEA platform of the Broad Institute (http://www.broadinstitute.org/gsea/index.jsp). For gene expression analysis in breast tumors, total RNA was extracted from frozen sections of breast tumor samples using the Nucleospin RNA set for Nucleozol kit according to the manufacturer’s protocol (Macherey-Nagel). Libraries were prepared using a modified version of the Takara SMARTer Stranded Total RNA-Seq Kit - Pico Input Mammalian kit (Hendricks et al., in preparation). In brief, 100 ng of RNA at 10 ng/μL was sonicated using RL230 Covaris sonicator (Covaris Inc) and resulting material was confirmed using a Fragment Analyzer (Agilent). 10 ng of each sonicated sample was prepared using the pico input kit as for FFPE samples using a 1:8 volume reduction on the STP MosquitoHV. Final libraries were validated by Fragment Analyzer and qPCR prior to sequencing on a NextSeq500 with 50nt single end reads. Reads were mapped to the human genome (USCS hg19 build) using TopHat v. 2.1.0 and transcript abundance was determined using HTSeqCount v. 0.9.1. Molecular subtyping, PAM50 and Claudin-low, was performed using the Genefu v.2.20.0 R package (http://www.pmgenomics.ca/bhklab/software/genefu). For gene expression analysis in mouse luminal cells and MaSC-basal cells, raw sequencing data for GSE60450 were downloaded from the GEO/SRA repository (ftp-trace.ncbi.nlm.nih.gov/sra) and reads were mapped to the UCSC mm9 mouse genome build (genome.ucsc.edu) using RSEM. Targeted pairwise differential expression analyses between basal and luminal samples were conducted using EBSeq v1.4.0 with median normalization. All RNA-seq analyses were conducted in the R Statistical Programming language. For gene expression analysis across breast tumor subtypes the pipeline from Fougner et al (*30*) was used on the TCGA-BRCA Dataset (github.com/clfougner/ClaudinLow). For ChIP-sequencing analysis, previously reported genome-wide murine Snail binding sites elucidated using ChIP-seq were obtained from the Gene Expression Omnibus (GSE61198). Mouse mm10 (NCBI Build 38/GRCm38 assembly) loci were translated to corresponding mm9 (NCBI Build 37 assembly) loci using the UCSC liftOver utility (command-line version timestamped 20170321). Peaks were annotated and labeled with corresponding genomic features using R package ChIPseeker (ver. 1.22.1) and UCSC mm9 knownGene annotation. Peaks within +/-3000bp of transcription sites were annotated as promoter-associated peaks. Mouse gene symbols were mapped to human equivalents using homology information from the Mouse Genome Informatics portal (MGI, informatics.jax.org) and promoter peaks from ChIP-seq analysis were subsequently associated with the 47 candidate ciliogenesis inducers along with differential gene expression results from Snail v. CTL RNA-seq analysis.

### Statistical analysis

Prism was used to analyze data, draw graphs and perform statistical analyses. Data are presented as the mean ± standard errors of mean (s.e.m.). Statistical analyses were carried out by Student’s t-test unless otherwise specified. * p ≤ 0.05, ** p ≤ 0.01, *** p ≤ 0.001 were considered significant.

## Data availability

RNA-sequencing data have been deposited in the Gene Expression Omnibus (GEO) under accession code GSE160549. Previously published data that were reanalyzed are available under the accession numbers GSE60450 and GSE61198.

## Supplementary Materials

Fig. S1: EMT program activation and ciliogenesis.

Fig. S2: EMT programs induce FGFR1 expression and FGF signaling controls MaSC stemness.

Fig. S3: GLIS2 inhibition occurs at the post-transcriptional level and *TRIM32* knockout does not affect the M-like status of HMLE cells.

Fig. S4: GLIS2Cter expression does not affect the M-like status of HMLE cells or primary ciliogenesis but affects cilium-signaling.

Fig. S5: EMT-dependent primary ciliogenesis results in GLIS2 inhibition in transformed cells and GLIS2Cter expression does not affect EMT and ciliogenesis but affects ciliary signaling.

Fig. S6: Claudin-low tumors express low levels of epithelial markers, display primary cilia, and express high levels of GLIS2 target genes.

Table S1: Snail direct target genes.

## General

We thank Xavier Pinson, Roselyne Viel and Aurore Nicolas for help in development of light sheet microscopy, immunohistochemistry methods, and processing of breast tumor samples, respectively; Julia Fröse for providing HMLE cells; Feng Zhang for providing the LentiCRISPR v2 plasmid (Addgene plasmid no. 52961); The Swanson and Biosit Biotechnology Centers, including MRic, H^2^P^2^, CRB Santé, Arche, the MIT BioMicro Center and Flow cytometry core facilities for technical support.

## Author contributions

M.M.W., C.C., J.A.L., V.J.G. designed research; M.M.W., C.C., S.M., M.L.G., A.S., Q.D., T.E.C., S.M., V.J.G. performed research, Q.D., F.G., P.T., A.M.J., J.E.V., R.A.W., M.A., C.P. contributed new reagents/analytic tools, M.M.W., C.C., M.L.G, A.S., A.B., J.A.L., V.J.G., analyzed data. M.M.W., J.A.L., V.J.G. wrote the paper.

## Data and materials availability

All data needed to evaluate the conclusions in the paper are present in the paper and/or the Supplementary Materials.

## Supplementary Materials

**Fig. S1:**
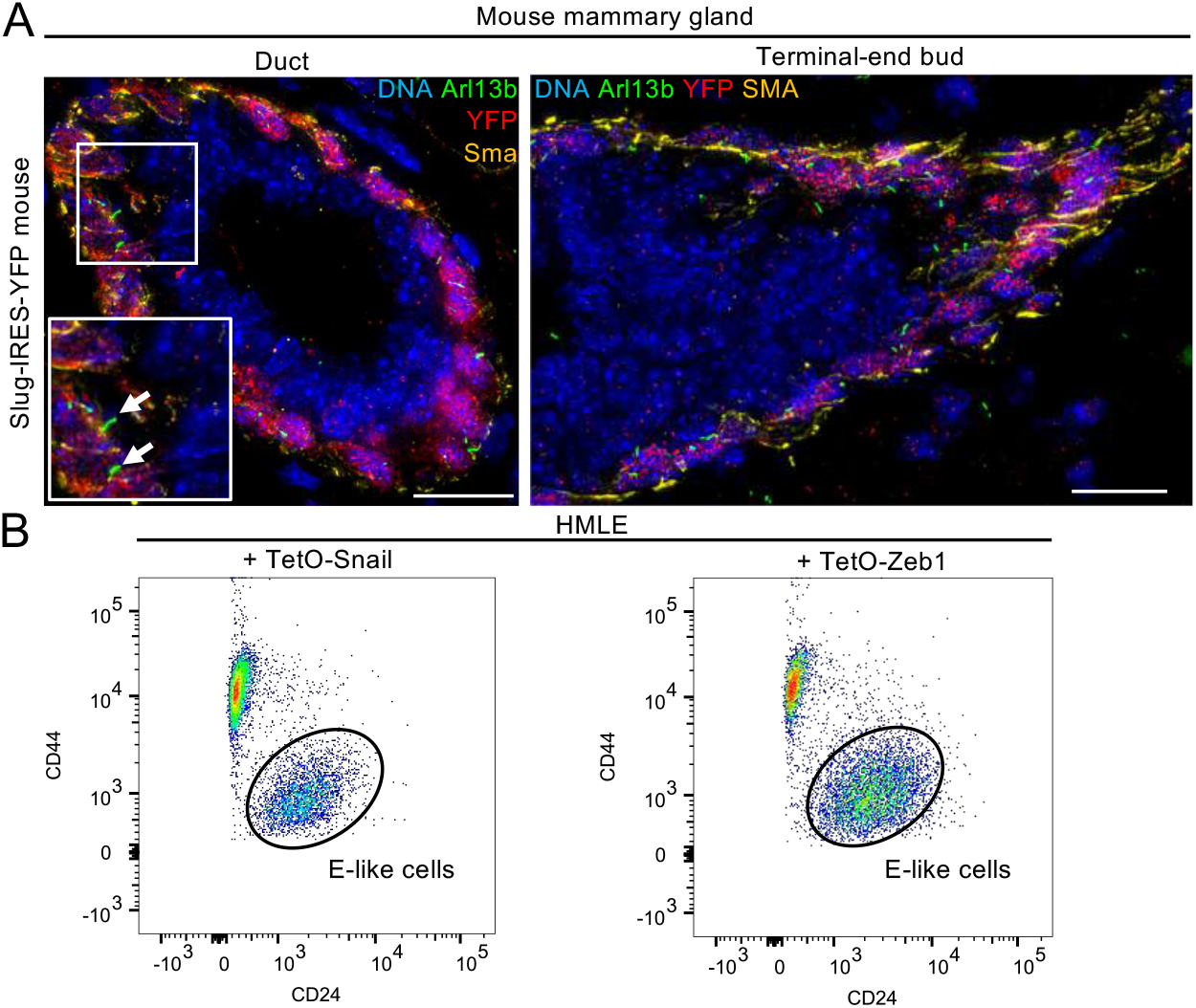
EMT program activation and ciliogenesis. (A) Mammary gland sections from Slug-IRES-YFP mice (8 weeks old) were stained for the indicated proteins. Representative images of a duct and of a remaining terminal-end bud (TEB) are shown. Scale bar: 15 μm (inset: 1.5X). (B) Epithelial-like (E-like, CD44^lo^ CD24^hi^) cells from the indicated un-induced HMLE variants were isolated by FACS using the indicated cell surface markers.

**Fig. S2:**
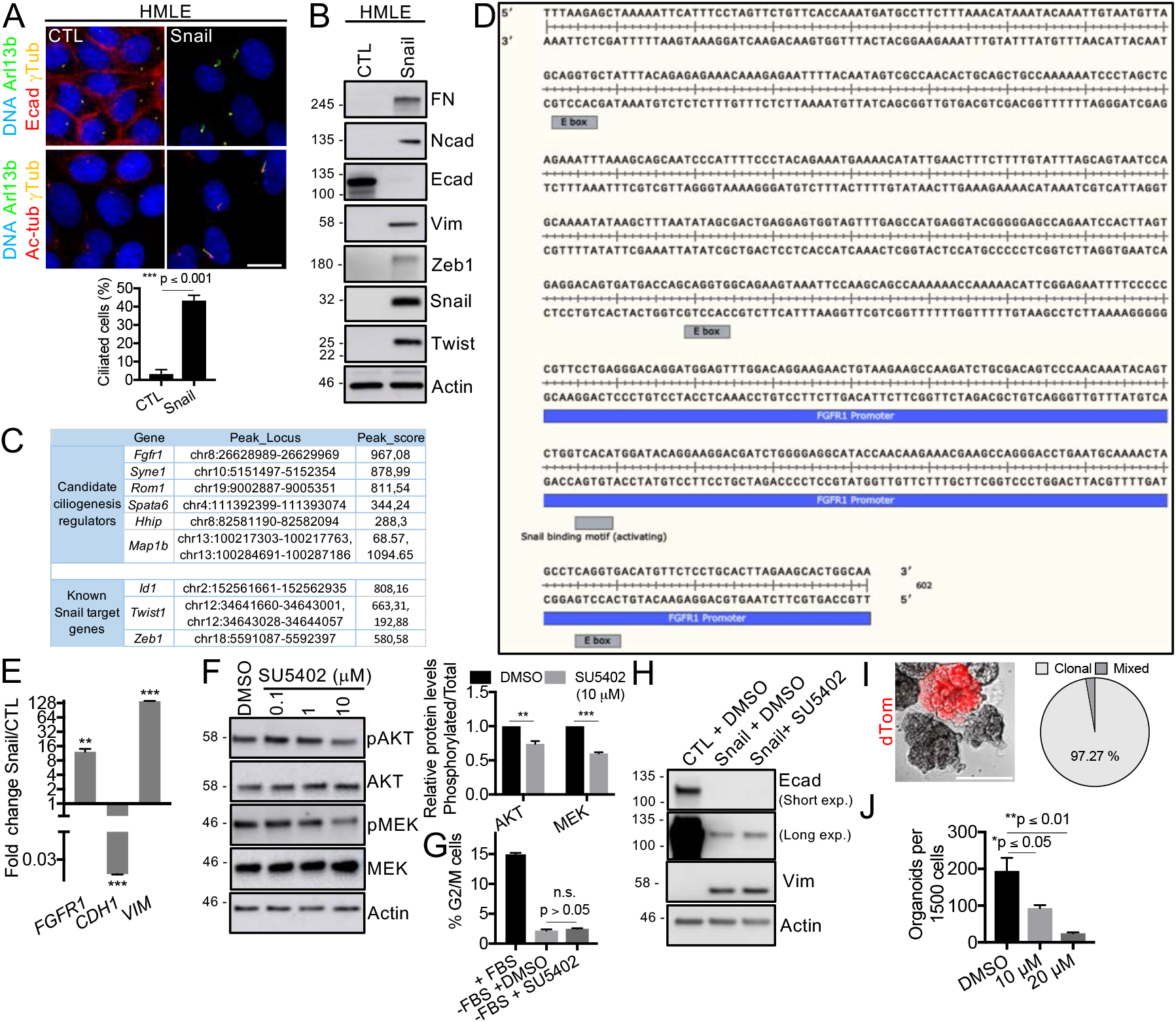
EMT programs induce FGFR1 expression and FGF signaling controls MaSC stemness. (A) Control (CTL) and Snail-expressing HMLE cells were grown until high-confluence, serum starved for 24 h, and stained for the indicated proteins. The percentage of ciliated cells was measured (n = 3, mean ± SEM). Representative results from 3 independent experiments are shown (Scale bar: 15 μm). (B) Western blot analysis of the protein expression levels of EMT markers in CTL or Snail-expressing cells. (C) Snail direct target genes were identified by CHIP-sequencing analysis using mammary tumor cells (GSE61198). Genes that were identified in Fig. 2C as putative ciliogenesis inducers were found in the list (*Fgfr1, Syne1, Rom1, Spata6*), along with known Snail direct targets (*Id1, Id2, Twist1, Zeb1, Vtn*). TSS/Peak coordinates are indicated for each gene, as well as the peak score, which is a normalized value of the read counts that were identified for each gene. (D) Snail-binding motifs associated to Snail dependent regulation of transcription (E-boxes or a binding motif associated to Snail-dependent activation of transcription^38^) were found in close proximity (within the 400 bp upstream) or in the Fgfr1 promoter (mm9 genome). (E) Relative levels of the indicated gene transcripts were determined by real-time qPCR analysis of CTL and Snail-expressing HMLE cells (n = 3, mean ± SEM). Representative results of 3 independent experiments are shown. (F) Western blot analysis of MEK and AKT phosphorylation in Snail-expressing HMLE cells treated with DMSO or SU5402 at the indicated concentrations. Phosphorylated and total protein levels were quantified (n = 3, mean ± SEM). Representative results of 3 independent experiments are shown. (G) Snail-expressing HMLE cells were exposed to serum or serum-starved with DMSO or SU5402 (10 μM). The percentage of cells in G2/M was determined by FACS analysis of the DNA content in each variant (n = 3, mean ± SEM). Representative results of 3 independent experiments. (H) Western blot analysis of the protein expression levels of EMT markers in CTL or Snail-expressing cells treated with DMSO or with SU5402 (10 μM). (I) MaSC-enriched basal cells that expressed dTomato (dTom) were mixed (1:1) with cells that did not express dTom and plated in 3D. The clonality of the organoid resulting from the co-culture was quantified by brightfield and fluorescence microscopy. Scale bar, 100 μm. (J) Organoid-forming capacity was determined for MaSC-enriched basal cells treated with SU5402 at the indicated concentrations 10 days after plating (n = 3, mean ± SEM). Representative results of 3 independent experiments. ** p ≤ 0.01, *** p ≤ 0.001.

**Fig. S3:**
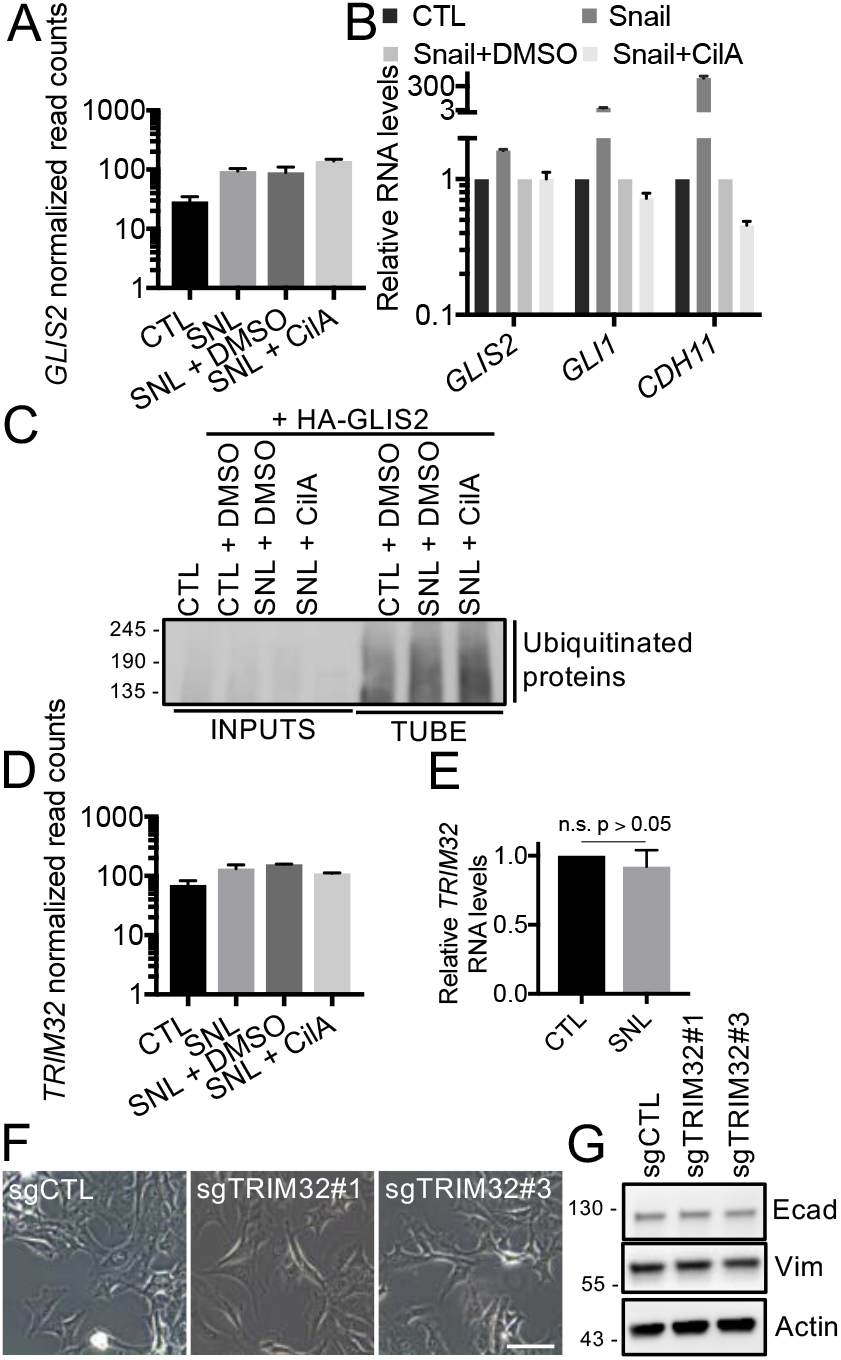
GLIS2 inhibition occurs at the post-transcriptional level and *TRIM32* knockout does not affect the M-like status of HMLE cells. (A) Expression of *GLIS2* was identified in the indicated HMLE variants by RNA-sequencing as described in the legend of Fig. 3 (mean ± SEM). (B)Relative levels of the indicated gene transcripts in the indicated cells were determined by real-time qPCR analysis (n = 3, mean ± SEM). Representative results of 3 independent experiments.(C)CTL and Snail-expressing (SNL) HMLE cells with or without HA-GLIS2 were treated or not with DMSO and Ciliobrevin A (CilA). Ubiquitinated proteins were purified by TUBE pull-down and analyzed by western blot. (D) Expression of *TRIM32* was identified by RNA-sequencing as described in the legend of Fig. 3 (mean ± SEM). (E) Relative levels of *TRIM32* were determined in the indicated cells by real-time qPCR analysis (n = 3, mean ± SEM). Representative results of 3 independent experiments. (F) The impact of *TRIM32* knockout in Snail-expressing HMLE cells on morphology was examined by brightfield microscopy. (G) The impact on expression of EMT markers was determined by western blot analysis.

**Fig. S4:**
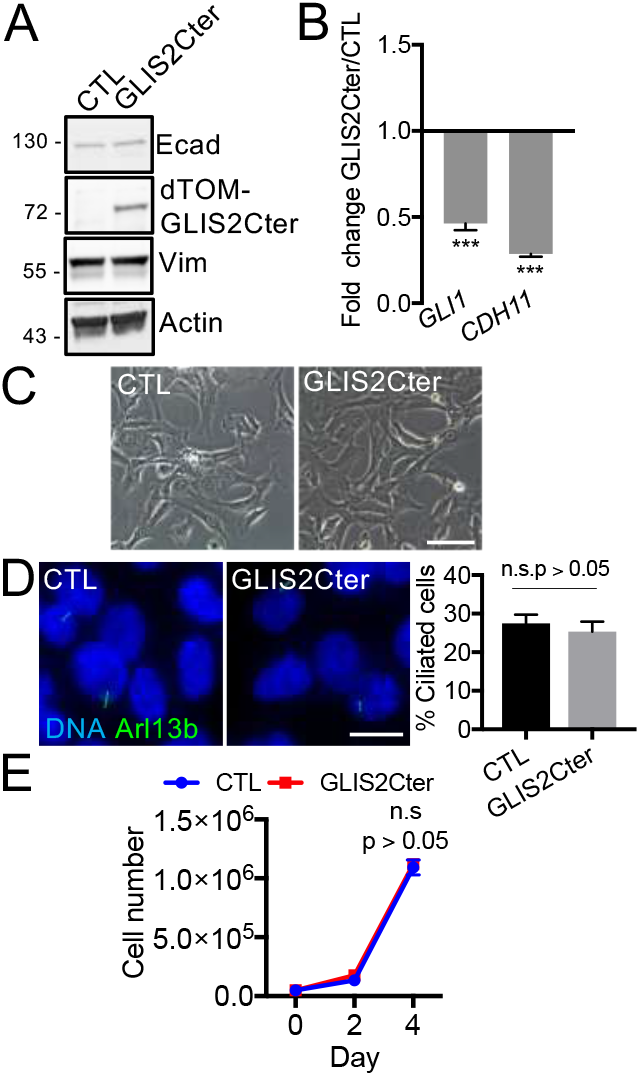
GLIS2Cter expression does not affect the M-like status of HMLE cells or primary ciliogenesis but affects cilium-signaling. (A) GLIS2Cter expression and its impact on expression of EMT markers in Snail-expressing HMLE cells were determined by western blot analysis. (B) Relative levels of *GLI1* and *CDH11* were determined in the indicated variants by real-time qPCR analysis (n = 3, mean ± SEM). Representative results of 3 independent experiments. (C-E) The impact of GLIS2Cter expression on morphology was examined by brightfield microscopy (C), on ciliogenesis was assessed by immunofluorescence microscopy (D), on proliferation in 2-dimension was analyzed by counting the total cell number per well (3 wells/line) at the indicated time points (E). All scale bars: Brightfield 100 μm, Immunofluorescence 15 μm.

**Fig. S5:**
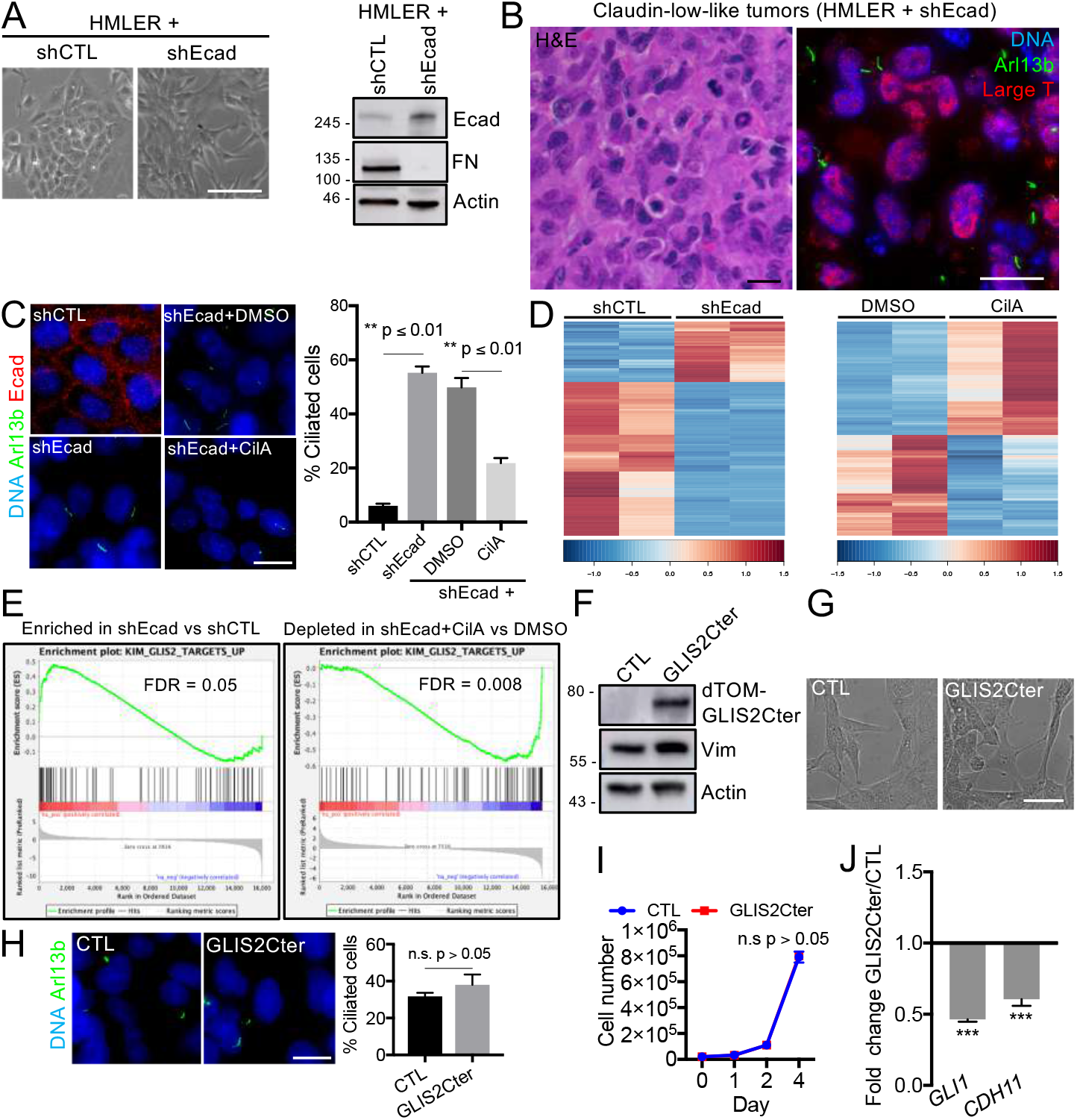
EMT-dependent primary ciliogenesis results in GLIS2 inhibition in transformed cells and GLIS2Cter expression does not affect EMT and ciliogenesis but affects ciliary signaling. (A) shControl (CTL)- and sh-Ecadherin (Ecad)-expressing HMLER cells were examined for morphology by brightfield microscopy (scale bar: 100 μm) and for their expression of EMT markers by western blot analysis. (B) Sections of tumors arising from HMLER + shEcad cells that were orthotopically transplanted in the fat pad of female mice, as described in the legend of Fig. 5, were stained with H&E as well as for the indicated proteins. Representative images are shown. Scale bars: 15 μm. (C, D) shCTL- and shEcad-expressing HMLE cells were grown until high confluence and serum starved for 24h without or with DMSO or Ciliobrevin A (CilA). (C) Cells were stained for the indicated proteins and the percentage of ciliated cells was determined (n = 3 mean ± SEM). Representative results from 3 independent experiments are shown. Scale bar: 15 μm. (D) Gene expression in the indicated HMLER variants was analyzed by RNA-sequencing. Heat-maps showing the differentially expressed genes (q-value ≤ 0.05, Fold-change ≥ 2) between samples are displayed. (E) A GSEA analysis shows significant enrichment of GLIS2 target genes in the list of upregulated genes in shEcad-versus (vs) shCTL-expressing cells (left panel) and depletion in the list of shEcad CilA-treated vs DMSO-treated cells (right panel) (False discovery rate (FDR) ≤0.05). (F-J) The impact of GLIS2Cter expression in HMLER + shEcad cells on expression of EMT markers and GLIS2 targets, on morphology, on ciliogenesis and cell proliferation was examined as described in the legend of Fig. S4. Scale bars: Brightfield 50 μm, Immunofluorescence 15 μm.

**Fig. S6:**
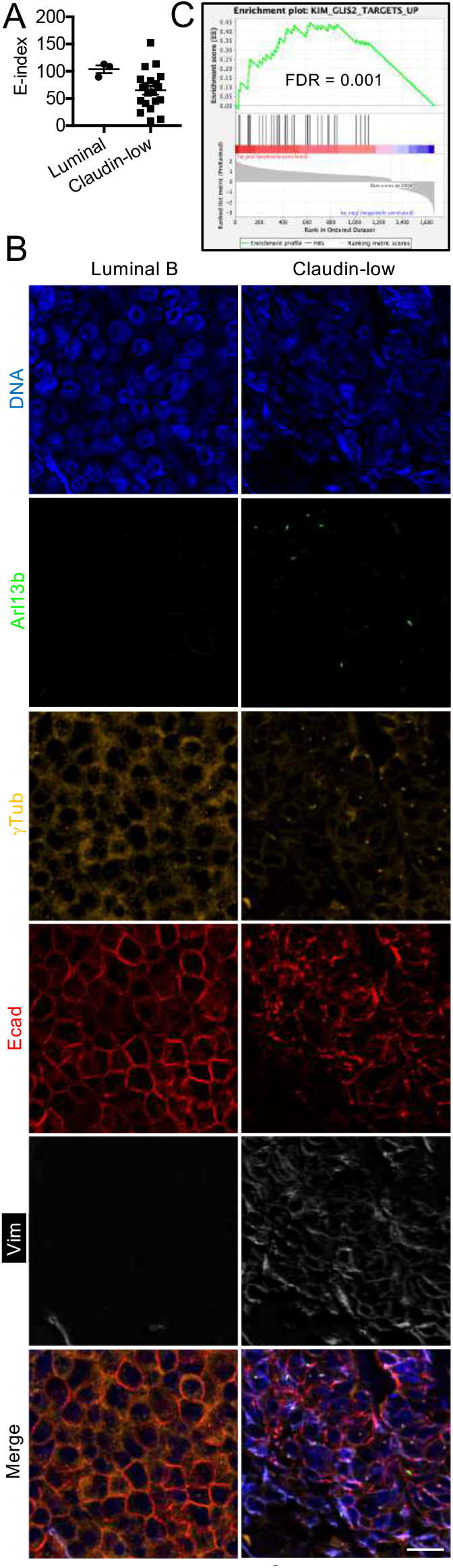
Claudin-low tumors express low levels of epithelial markers, display primary cilia, and express high levels of GLIS2 target genes. (A-B) Paraffin sections of breast tumors were stained as described in the legend of Fig. 6B-C. (A) Ecadherin, Claudin 4 and Claudin 7 signal intensities were measured and used to define an Epithelial-index (E-index = (Ecadherin signal intensity + Claudin 4 signal intensity + Claudin 7 signal intensity)/3). (B) Scale bar: 20 μm. (C) A GSEA analysis shows significant enrichment of GLIS2 target genes (FDR<0.05) in the list of upregulated genes in claudin-low cancers versus other breast cancers defined by Prat et al^29^.

**Table S1:**
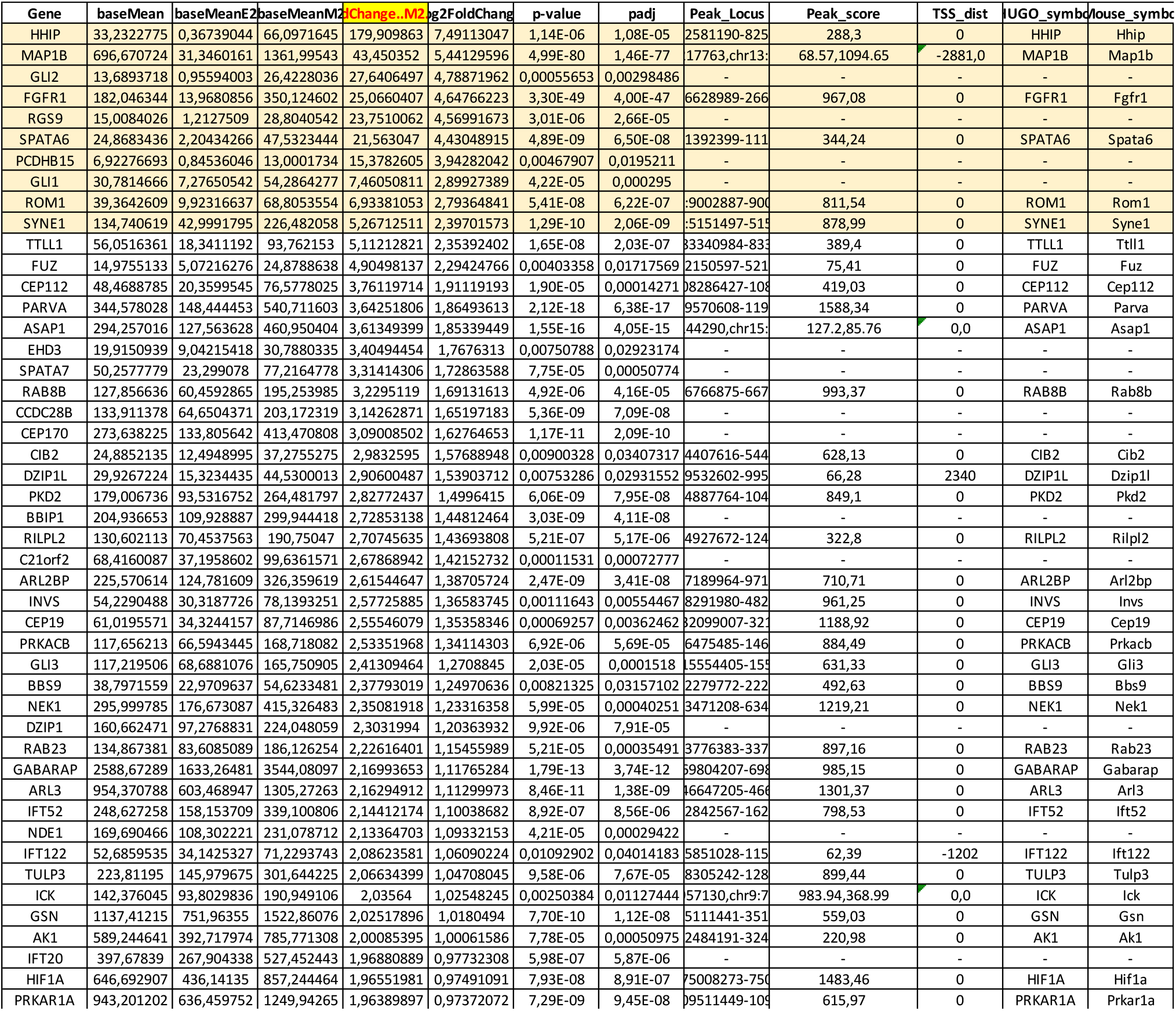
Snail direct target genes. Candidate ciliogenesis inducers with Snail v. CTL RNA-seq differential gene expression results and associated promoter peaks (+/- 3kbp) from Ye et al. GSE61198 (baseMean = Mean gene expression across samples; padj = adjusted p-value; TSS_dist = Distance of peak from transcription start site of gene; HUGO = Human Genome Organization/Human Gene Nomenclature Committee).

